# DUF581-9 (At2g44670; FLZ3) negatively regulates SnRK1 activity by interference with T-loop phosphorylation

**DOI:** 10.1101/2022.03.17.484690

**Authors:** Jennifer Bortlik, Jost Lühle, Saleh Alseekh, Christoph Weiste, Alisdair R. Fernie, Wolfgang Dröge-Laser, Frederik Börnke

## Abstract

In plants, SUCROSE NON-FERMENTING RELATED KINASE 1 (SnRK1) is a key energy-sensor that orchestrates large-scale transcriptional reprogramming to maintain cellular homeostasis under energy deficit. SnRK1 activity is under tight negative control, although the exact mechanisms leading to its activation are far from being understood. We show that the Arabidopsis DOMAIN OF UNKNOWN FUNCTION (DUF581) protein DUF581-9/FLZ3 binds to the catalytic SnRK1α subunit KIN10 to inhibit its activation by GRIK-dependent T-loop phosphorylation. Overexpression of *DUF581-9* in Arabidopsis dampens SnRK1 signalling and interferes with adaptation to dark-induced starvation. The presence of DUF581-9 significantly reduced SnRK1 activity in protoplasts and *in vitro*. This was accompanied by a reduction in T175 T-loop phosphorylation and also diminished KIN10 auto-phosphorylation. Furthermore, DUF581-9 reduced binding of the up-stream activating kinase GRIK2 to KIN10, explaining the reduced KIN10 T-loop phosphorylation. Ectopically expressed DUF581-9 protein was rapidly turned over by the proteasome when Arabidopsis plants were subjected to starvation treatment, likely releasing its inhibitory activity on the SnRK1 complex. Taken together, our results support a model in which DUF581-9 negatively regulates SnRK1 activity under energy sufficient conditions. Turnover over of the protein provides a rapid way for SnRK1 activation under energy deficit without the need of de-novo protein synthesis.

## Introduction

In plants, biotic and abiotic stresses typically lead to cellular energy depletion. As a response, transcriptional and metabolic reprogramming of the cell results in the downregulation of energy-consuming processes, including a cessation of growth, and an induction of catabolic reactions to ensure resource allocation in support for stress tolerance and survival (Baena-Gonzalez, 2010). A central regulator of the low-energy response is the energy-sensor protein kinase Sucrose Non-Fermenting related kinase 1 (SnRK1), which is orthologous to the AMP-dependent kinase (AMPK) and Sucrose Non-Fermenting 1 (SNF1) in mammals and yeast, respectively (Broeckx *et al*., 2016). Stress-mediated activation of SnRK1 results in the direct phosphorylation of several central biosynthetic enzymes to reduce their activity (Halford &Hey, 2009; Nukarinen *et al*., 2016). In addition, SnRK1 phosphorylates a range of different transcription factors (summarized in Broeckx *et al*., 2016) to mediate large scale transcriptional reprogramming of the cell. Transient overexpression of KIN10 in Arabidopsis protoplasts resulted in a transcriptional profile reminiscent of various starvation conditions and led to the identification of approximately 1.000 putative SnRK1 target genes (Baena-González *et al*., 2007).

The SnRK1 holoenzyme is a heterotrimeric complex consisting of a catalytic α subunit and regulatory β- and γ-subunits (Broeckx *et al*., 2016). The α subunit consists of a highly conserved N-terminal Ser/Thr kinase domain and a C-terminal regulatory domain, which mediates interaction with the β- and γ -subunits. The β-subunit acts as complex scaffold but also contributes to complex localization and substrate specificity (Ramon *et al*., 2019). In Arabidopsis (*Arabidopsis thaliana*), the catalytic α subunit is represented by two isoforms, KIN10 (At3g01090) and KIN11 (At3g29160), both of which are expressed ubiquitously, although KIN10 accounts for the majority of SnRK1 activity (Jossier *et al*., 2009). SnRK1 activity is regulated by phosphorylation/dephosphorylation of a T-loop threonine in the catalytic α subunit (T175 of AKIN10 and T176 in AKIN11) involving upstream kinases GRIK1/2 (Shen *et al*., 2009). SnRK1 also undergoes significant auto-phosphorylation (Baena-González *et al*., 2007), and a model has been put forward in which upstream kinases are required for initial phosphorylation and activation of newly synthesized SnRK1 proteins, while full activation requires additional self-phosphorylation events. While for SNF1 from yeast and AMPK from mammals a strict correlation between T-loop phosphorylation with kinase activity on the one hand, and cellular energy level and T-loop phosphorylation on the other hand occurs, this correlation is much less clear for SnRK1. Although SnRK1 T-loop phosphorylation appears to be required for kinase activity *per se* (Glab *et al*., 2017), its overall phosphorylation level does not change substantially even under conditions in which there is either an increase or decrease in SnRK1 activity (Baena-González *et al*., 2007; Coello *et al*., 2012; Rodrigues *et al*., 2013).

Recent evidence suggests that, rather than being activated upon energy deficit like AMPK and SNF1, SnRK1 is activated by default but its activity is repressed under energy sufficient conditions (Crepin &Rolland, 2019; Ramon *et al*., 2019). Several molecules, including metabolites and proteins, have been described to negatively regulate SnRK1 signalling. For instance, the intermediate of trehalose biosynthesis, trehalose-6-phosphate (T6P), which acts as fuel gauge to signal sucrose availability (Lunn *et al*., 2006), was shown to inhibit SnRK1 activity in a tissue- and developmental stage-specific manner (Zhang *et al*., 2009; Debast *et al*., 2011; Martínez-Barajas *et al*., 2011). Recently, T6P was shown to directly bind to the SnRK1 α subunit to weaken the GRIK1-KIN10 association and thus interfering with T-loop phosphorylation (Zhai *et al*., 2018). Emanuelle et al., (2015) identified a heat-labile, >30 kDa, soluble proteinaceous factor present in the lysate of young, growing, 3 week old rosette leaves that significantly inhibited SnRK1 catalytic activity when added to the recombinant kinase complex *in vitro*. However, the identity of this factor let alone its mode of action is currently unknown. Known SnRK1 interacting proteins that negatively regulate its signalling activity include PLEIOTROPIC REGULATORY LOCUS1 (Bhalerao *et al*., 1999) and SnRK1A INTERACTING NEGATIVE REGULATOR PROTEINS (SKINs). The latter interact with catalytic domain of SnRK1A in rice in order to antagonise SnRK1 activity and thus prevent its over-activation in response to abscisic acid signalling (Lin *et al*., 2014).

We have previously demonstrated that proteins containing a DOMAIN OF UNKNOWN FUNCTION (DUF) 581 interact with the catalytic α subunits of SnRK1 (AKIN10/11) from Arabidopsis via their zinc-finger containing DUF581 domain (Nietzsche *et al*., 2014). DUF581 proteins are confined to the plant kingdom and constitute a family of 18 members in Arabidopsis. Besides the variable N- and C-termini, the conserved DUF581 is necessary and sufficient for the proteins to bind to KIN10/11, indicating that the variable portion could impart some sort of functional specificity to individual isoforms. Expression of the DUF581 genes in Arabidopsis is highly responsive to hormones and environmental cues, such as ABA treatment, hypoxia, heat, and nutrients (Nietzsche *et al*., 2014; Jamsheer &Laxmi, 2015). In addition to SnRK1, DUF581 proteins interact with a range of additional proteins many of which participate in central cellular signalling pathways or act in transcriptional regulation (Nietzsche *et al*., 2016). However, a functional relationship between SnRK1 signalling and DUF581 proteins has so far not been firmly established. A recent study suggests that at least two of the DUF581 proteins might act as negative regulators of SnRK1 activity, although their mode of action remains unclear (Jamsheer *et al*., 2018).

In the present study, we demonstrate that the DUF581-family member DUF581-9 (At2g44670/FLZ3) acts as a negative regulator of SnRK1 *in planta* and *in vitro*. Overexpression of *DUF581-9* in transgenic Arabidopsis lines attenuates SnRK1-mediated responses. We show that DUF581-9 reduces SnRK1 activity by binding to KIN10 and weakening its interaction with the upstream activating kinase GRIK. The resulting reduction in T-loop phosphorylation prevents further auto-phosphorylation events. DUF581-9 itself appears to be degraded by the proteasome under energy limiting conditions, thus relieving its inhibitory effect on the signalling pathway when required.

## Materials and Methods

### Plant material

*Arabidopsis thaliana* plants were grown on soil in an 8 h light / 16 h dark cycle (22°C / 18°C), 70 μmol m^− 2^ s^− 1^ light intensity and 70 % humidity. Transgenic plants were generated in the Col-0 ecotype. T-DNA insertion line *duf581-9 ko* (SALK_062585) was obtained from Nottingham Arabidopsis Stock Centre (NASC). Transgenic overexpression lines *DUF581-9 OE #1* and *#2* were generated following the *Agrobacterium tumefaciens* (strain GV3101)-based transformation via the floral dip method (Clough &Bent, 1998), and independent transformants were selected by Kanamycin selection. *Nicotiana benthamiana* wildtype plants were grown on soil with a 16 h light / 8 h dark cycle.

### Plasmid construction

Full-length coding regions of DUF581-9/ FLZ3 (At2g44670) and of the DUF581-9_C47S_ variant (Nietzsche *et al*., 2014) were cloned into pENTR-D/TOPO (Thermo Fisher). For stable overexpression lines, the vector pRB-35S-3xmyc (Bartetzko *et al*., 2009) was used. For transient expression in *N. benthamiana*, the DUF581-9 sequence was inserted into pRB-35S-mCherry. Constructs for Bi-molecular Fluorescence Complementation analysis are based on Gateway®-cloning (GW) compatible versions of pRB-C-Venus^N173^ and pRB-C-Venus^C155^ (Üstün &Börnke, 2015). To create the *Pro*_*DUF581-9*_*::GUS* construct, a 1828 bp fragment upstream of the At2g44670 gene was amplified and inserted into the GW compatible vector pBGWFS7,0 (Karimi *et al*., 2002). For recombinant protein expression we used GW versions of pMAL-C2 (New England Biolabs) and pDEST-17 (Thermo Fisher) with N-terminal MBP-or GST tag, respectively. For yeast two-hybrid analyses, fragments were recombined into GW versions of the GAL4 DNA-binding domain vector pGBT-9 and the activation domain vector pGAD424 (Clontech). Oligonucleotides used for cloning are listed in Supplementary Table S3.

### Starvation and survival assays

Short term starvation experiments were conducted with 4 week old plants, grown under SD conditions as stated above. Thirty minutes into the photoperiod, lights were switched off for 6 h before samples were taken. To analyse the survival rate after long term starvation, 3 week old Arabidopsis plants grown under SD conditions were subjected to a dark-treatment for 10 days. Recovery performance was quantified 7 days after resuming normal light and growth conditions. Plants with a clearly visible intact green meristem were considered survivors.

### Analysis of GUS reporter lines

Different developmental stages of several independent transgenic *Pro*_*DUF581-9*_*::GUS* lines were investigated for tissue specific localisation patterns using histochemical staining according to Jefferson et al. (1987).

### RNA extraction and qRT-PCR

RNA was extracted from ground leaf tissue using NucleoZOL (Machery-Nagel) reagent. First strand cDNA synthesis and quantitative real-time RT-PCR (qRT-PCR) was conducted as described (Nietzsche *et al*., 2018). Oligonucleotides used for qRT-PCR are listed in Supplementary Table S3.

### Immuno-affinity purification

Two grams of frozen Arabidopsis leaf material was ground in liquid nitrogen and thawed in 4 ml extraction buffer (100 mM Tris, pH 8.0; 100 mM NaCl; 5 mM EDTA; 5 mM EGTA; 20 Mm DTT; 10 mM NaF; 10 mM Na_3_VO_4_; 2 µl/ml plant protease inhibitor cocktail [Sigma P9599]; 0.5 % [v/v] Triton X-100). After centrifugation, the supernatant was incubated with 40 μL of magnetic Myc-Trap® (50 % [v/v] slurry, Chromotek) and incubated for 60 min at 4°C. After five washing steps (100 mM Tris, pH 8.0; 100 mM NaCl; 0.5 mM EDTA; 1 mM DTT) the purified myc-tagged DUF581-9 protein was either used for immunoblot or for LC-MS/MS analysis.

### Immunoblotting

Protein extracts were boiled with 5 x SDS sample buffer and separated by SDS-page. Immunoblotting was carried out with α-myc antibody (1:2.500, Abcam), followed by Goat Anti-Rabbit IgA alpha chain (HRP) (1:5.000) secondary antibody (Abcam). KIN10 was detected by an α-KIN10 (1:500) antibody (Agrisera). To detect KIN10 T-loop phosphorylation, a phospho-AMPK alpha-1 (Thr172) polyclonal antibody (Sigma) was used (1:500). Signals were visualized using chemiluminescence (Thermo Fisher Scientific) with a ChemiDoc Imaging system (Biorad).

### Purification of recombinant proteins

Recombinant proteins were expressed either in *E. coli* M15 (MBP-fusions) or *E. coli* BL12 (GST-fusions) cells. Bacteria were lysed by sonication. After centrifugation, the MBP-fusion proteins were purified using amylose resin (New England Biolabs) and GST-fusions by glutathione-resin (GE Healthcare), each according to the manufacturer’s instructions.

### Pull-down assays

Purified KIN10-GST recombinant protein was incubated with crude extract of MBP EV, MBP-GRIK2, MBP-DUF581-9 or the C47S variant in pull-down buffer (20 mM Tris-HCl, pH 8; 100 mM NaCl; 1 mM EDTA) at 4 °C for 1 h. Subsequently, the gluthatione-resin was washed five times with pull-down buffer. The proteins were eluted from beads by boiling in 80 μl 2 × SDS sample buffer and separated on 12 % SDS-PAGE gels. Gel blots were analysed using anti-MBP (NEB, 1:10.000) and anti-GST antibodies (Santa Cruz Biotechnology, 1:1.000).

### In vitro SnRK1 kinase assay

*In vitro* kinase assay was performed with purified MBP-tagged *At*KIN10, *At*KIN10_K48R_, *At*GRIK2, *At*DUF581-9 and *At*DUF581-9_C47S_ proteins as well as MBP-SAMS peptide expressed in *E*.*coli*. The reactions were started by adding kinase buffer (50 mM Tris HCl, pH 7.5; 10 mM MgCl_2_; 0.1 mM EGTA; 1 mM DTT; 0.1 mM ATP), subsequently incubated for 30 min at 30°C and stopped by adding SDS sample buffer or directly frozen in N_2_ for further LC-MS/MS analysis. Alternatively, the assay was carried out in the presence of 2 µCi _32_P γ-ATP and the incorporation of radiolabel into the SAMS peptide was carried out as described previously (Debast *et al*., 2011).

### Protoplast isolation and transformation

Protoplast transfection assays were performed according to Yoo et al. (2007) with modifications as described in (Ehlert *et al*., 2006). GUS enzyme assays were performed after 16 h of incubation in light (110 µmols^−1^m^−2^). Reporter and effector plasmids have previously been described (Pedrotti *et al*., 2018; Henninger *et al*., 2022).

### Mass Spectrometry and Data Analysis

Bead-bound DUF581-9-myc was supplemented with 0.1 % RapiGest SF (Waters, Eschborn, Germany), whereas *in vitro* kinase assay mixture was directly reduced and digested following the method of Kaspar et al. (2010). Subsequently, desalting of peptides was carried out according to Witzel et al. (2019). Protein digest were analysed using the Thermo Fisher Q Exactive high field mass spectrometer by reverse-phase HPLC-ESI-MS/MS using the Dionex UltiMate 3000 RSLC nano System coupled to the Q Exactive High Field (HF) Hybrid Quadrupole Orbitrap MS (Thermo Fisher Scientific) and a Nano-electrospray Flex ion source (Thermo Fisher Scientific) as described previously by Witzel et al. (2019). Each sample was measured in triplicate. Proteome discoverer software (PD2.4) was used to analyse and align the LC-MS raw data files, with its built-in MS Amanda, MS Mascot and Sequest HT search engine (Thermo Scientific) (Witzel *et al*., 2019). The MS/MS spectra were searched against Swissprot database for *A. thaliana* (UP000006548) and common contaminants for protein identification. Analysis parameter were set as described by Witzel et al. (2019). The result lists were filtered for high confident peptides and their signals were mapped across all LC-MS experiments (Col-0, *DUF581-9 #1, DUF581-9 #2* lines) and normalised to the total peptide amount per same LC-MS/MS experiment. Only unique peptides were selected for quantification and abundances of all peptides allocated to a specific protein were summed and compared.

### Metabolite analysis

4 week old Arabidopsis plants grown under SD conditions were treated as described above. Rosettes of 6 biological replicates were used for metabolite profiling and performed exactly as described by Lisec et al. (2011). Metabolite identities were verified via comparison to spectral libraries of authentic standards housed in the Golm Metabolome Database (Kopka *et al*., 2005).

### Yeast two-hybrid (Y2H) analyses

Direct protein-protein interaction was tested by Y2H technique according to the yeast protocols handbook and the Matchmaker GAL4 Two-hybrid System 3 manual (both Clontech, Heidelberg, Germany). Yeast strain Y190 was co-transformed with respective plasmids, followed by selection of transformants on medium lacking Leu and Trp at 30°C for 3 days and subsequent transfer to medium lacking Leu, Trp and His (supplemented with 25 mM 3-amino-triazole) for growth selection. Cells growing on selective medium were further tested for activity of the *lacZ* reporter gene using filter lift assays.

### Bimolecular fluorescence complementation (BiFC) analysis

To investigate *in planta* protein-protein interaction between KIN10-Venus^N173^ and GRIK2-Venus^C155^ in absence / presence of DUF581-9 mCherry, constructs were transformed into *A. tumefaciens* GV3101 and transiently expressed by Agrobacterium-infiltration in *N. benthamiana*. Cytosolic FBPase (At1g43670) served as positive control (Üstün &Börnke, 2015), whereas SPP2 mCherry (At2g35840) and FBPase mCherry were co-infiltrated as negative control. The BiFC-induced YFP fluorescence was detected by CLSM (LSM880, Axio Observer; Zeiss) after 48 hpi. The specimens were examined using the LD LCI Plan-Apochromat 25x/0.8 water-immersion Imm Korr DIC M27 objective for detailed images with excitation using the argon laser (YFP – 514 nm, mCherry - 561nm). Fluorescence was quantified by ImageJ (Fiji).

## Results

### DUF581-9 expression is down-regulated during dark-induced starvation

In order to obtain a deeper understanding concerning the functional relevance of the DUF581/KIN interaction for SnRK1 signalling, we initially concentrated our efforts on DUF581-9 (FLZ3; At2g44670), the smallest member of the protein family with only 9 amino acid residues N-terminal and 34 residues C-terminal, respectively, of the DUF581. To analyse the expression of *DUF581-9* under SnRK1 activating conditions, 4 week old Arabidopsis plants were subjected to sudden darkness or hypoxia for 24 h and subsequently mRNA levels were monitored by qRT-PCR. As shown in Fig. 1, *DUF581-9* expression was reduced by approximately 50 % in dark treated rosette leaves as compared to the control. Conversely, when dark treated plants were allowed to recover in SD for 19 h, *DUF581-9* expression was induced approximately 2-fold relative to untreated leaves. Submergence treatment did not affect *DUF581-9* expression (Fig. 1). Thus, *DUF581-9* expression is reduced in some but not all conditions associated with the activation of SnRK1 signalling while an induction of expression occurs during the recovery from starvation.

**Fig. 1.**
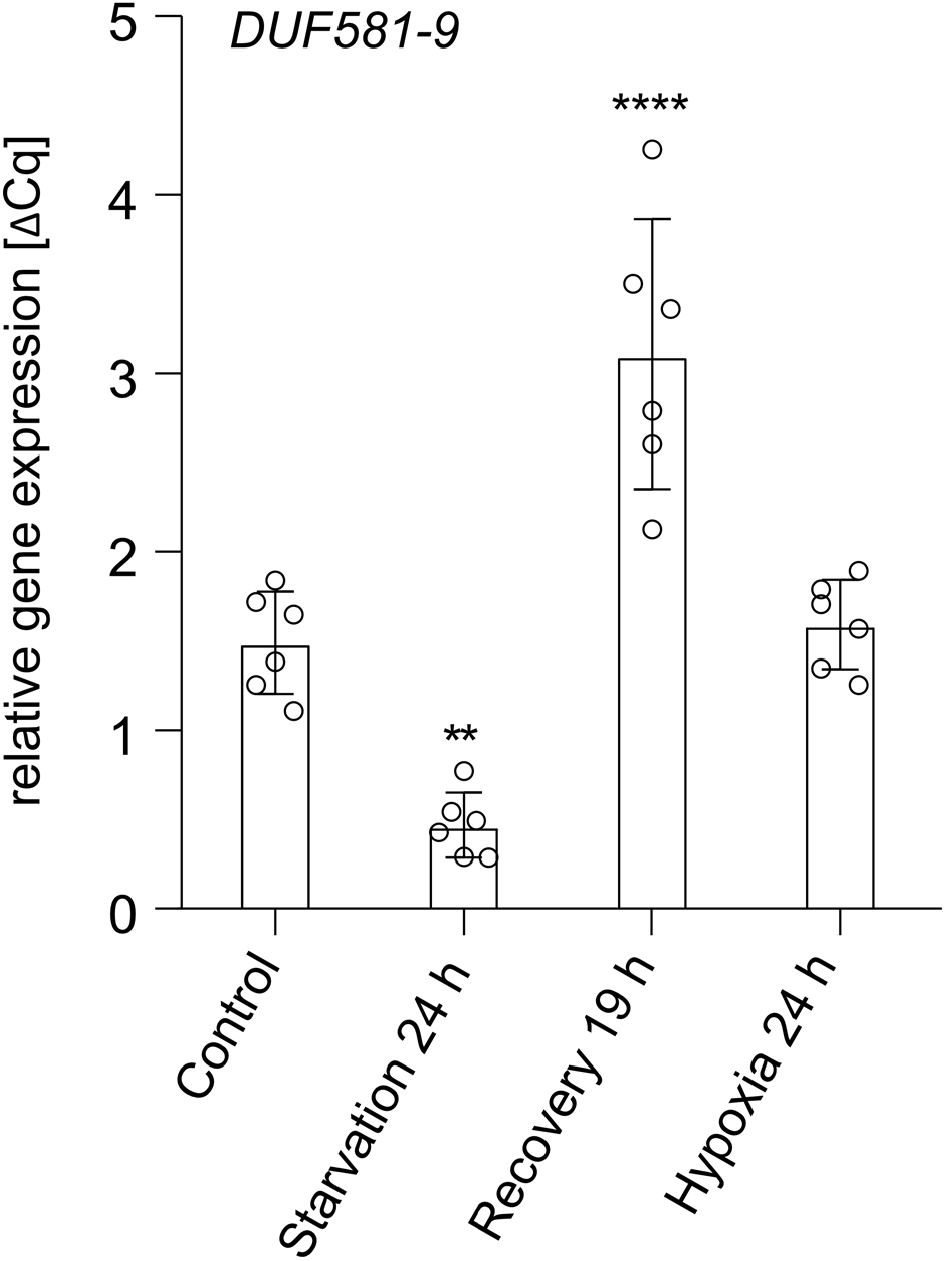
*DUF581-9* expression is reduced under starvation conditions. 4 week old Col-0 Arabidopsis plants grown in short day (8h light/16 h dark) were subjected to 24 h darkness followed by 19 h recovery under normal light conditions. For hypoxia treatment, plants were submersed in water for 24 h. Whole rosettes were harvested and used for RNA extraction and subsequent cDNA synthesis followed by qRT-PCR. *UBIQUITIN CONJUGATING ENZYME 9* (*UBC9*) was used as reference gene. Error bars indicate standard deviation (SD). Asterisks indicate statistical significant differences relative to the untreated control (ANOVA analysis followed by Sidak’s multiple comparison test) \*\**P* < 0.01, \*\*\*\**P* < 0.0001. The experiment was repeated twice with similar results.

To investigate *DUF581-9* expression at a higher spatial resolution, a *Pro*_*DUF581-9*_*::GUS* fusion construct was transformed into Arabidopsis. Multiple transgenic lines were examined for *GUS* activity at various developmental stages (Supporting Information Fig. S1). In vegetatively growing plants, strongest GUS staining was observed in and around the shoot apical meristem and in the proximal regions of the young emerging true leaves (Fig. S1a). As the leaves grew, *GUS* expression decreased in a tip-to-base manner. No *GUS* expression could be observed in mature leaves or other tissues of vegetatively growing plants over the observation period of 35 days post germination. In accordance with the qRT-PCR data of *DUF581-9* expression, GUS staining faded when the plants were subjected to dark induced starvation and resumed during a recovery period (Fig. S1a). In flowers of *Pro*_*DUF581-9*_*::GUS* lines staining was mainly confined to the pistil (Fig. S1b).

### Modulation of DUF581-9 expression affects SnRK1-dependent responses in Arabidopsis

To study potential functional links between DUF581-9 and SnRK1 signalling *in planta*, a *duf581-9* knock-out line as well as two transgenic Arabidopsis lines constitutively overexpressing *DUF581-9* were established (*DUF581-9 OE #1 and #2*) (Fig. S2). Lines with altered *DUF581-9* expression were grown alongside with Col-0 control plants under SD conditions in soil for 4 week. The expression of selected SnRK1 marker genes (Baena-González *et al*., 2007) was investigated by qRT-PCR. As previously reported, 6 h of starvation treatment led to a strong induction of *DARK INDUCED 6* (*DIN6*), encoding ASPARAGINE SYNTHETASE 1, as well as of *TREHALOSE-6-PHOSPHATE SYNTHASE 8* (*TPS8*) in control plants (Fig. 2a). In the case of *DIN6* this induction was significantly enhanced in *duf581-9* plants, while for both SnRK1 marker genes expression was reduced in *DUF581-9* OE lines as compared to the control (Fig. 2a). In order to investigate if modulation of *DUF581-9* expression affects the ability to cope with carbon limitation, wild type, *DUF581-9* OE and knock-out plants were grown for 3 week under SD conditions. Subsequently, the plants were subjected to a dark period of 10 days followed by a recovery for further 7 days under the initial growth conditions. Plant survival was assessed based on the emergence of new leaves during the recovery period. Approximately 80 % of the wild type control plants were able to survive the treatment while rate was reduced to 40 % and 20 % in line *DUF581-9 OE #1 and #2*, respectively (Figs. 2b and S3). The *duf581-9* line displayed unaltered survival relative to control plants (Figs. 2b and S3).

**Fig. 2.**
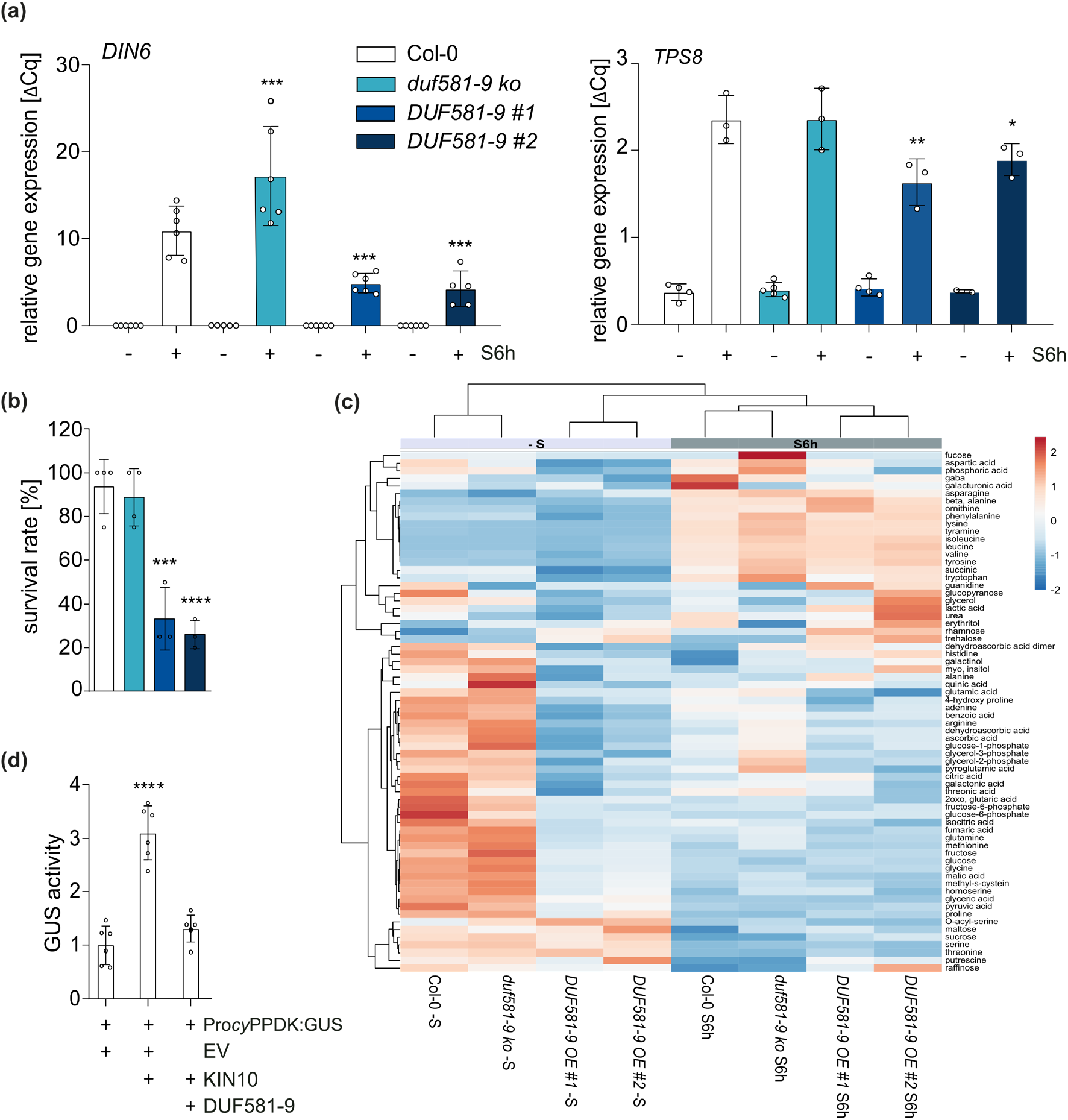
Ectopic overexpression of *DUF581-9* inhibits SnRK1 signalling and decreases plant survival under starvation. (a) Expression of SnRK1 marker gene *DARK INDUCED 6* (*DIN6)* and *TREHALOSE-PHOSPHATE SYNTHASE 8 (TPS8)* in 4 week old plants exposed to 6 h darkness (S6h). *UBC9* was used as reference gene. Error bars indicate SD. Asterisks indicated statistical significant differences relative to the Col-0 control (ANOVA analysis followed by Sidak’s multiple comparison test) \**P* < 0.05, \*\**P* < 0.01, \*\*\**P* < 0.001. The experiment was repeated at least three times. (b) Survival rate was determined in percentage after 10 days of dark-induced starvation and 7 days recovery of 4 week old Col-0, *duf581-9 ko* and *DUF581-9 OE* plants. Error bars indicate SD. Statistical analysis was carried out using ANOVA followed by Dunnett’s post-test. Significant differences against Col-0 ***P < 0.001, ****P < 0.0001. Three repetitions showed similar results. (c) GC-MS analysis of 4 week old Col-0, *duf581-9 ko* and *DUF581-9 OE* plants (n=6). Plants were grown in soil under short-day conditions (8 h of light / 16 h of dark), and rosettes were harvested either 6 h after the onset of light or after 6 h of darkness (30 min ZT). Metabolite data represent absolute values during light and dark conditions. The log2 fold change scale is indicated above the heat map. A negative score, depicted in blue, represents decreased levels, while a positive score (red colour) shows increased levels. The depth of the colour corresponds to the magnitude of the change in levels. Rows are centred; unit variance scaling is applied to rows. Rows are clustered using correlation distance and average linkage. Columns are clustered using Euclidean distance and average linkage. For details, see Supplemental Table S1. The experiment was repeated twice with comparable results. (d) Investigation of *in vivo* SnRK1 activity in a protoplast transactivation assay based on the SnRK1 responsive promoter of *cyPPDK2* fused to the GUS reporter gene. The Pro*cyPPDK::GUS* reporter construct was co-expressed with CaMV35S driven *KIN10* or *DUF581-9* effector constructs, respectively. Error bars indicate SD (n=6). Asterisks mark statistical significant differences relative to the EV control (ANOVA analysis followed by Dunnett’s multiple comparison test) **** = *P* < 0.0001. The experiment was carried out twice with comparable results.

Metabolite profiling of Arabidopsis plants with altered expression of DUF581-9 revealed major metabolic shifts in *DUF581-9 OE* lines. While the *duf581-9* mutant clustered with control plants, *DUF581-9 OE* lines formed a separate cluster in light grown plants as well as after 6 h starvation treatment (Fig. 2c; Table S1). During the light period, a few metabolites, including minor carbohydrates such as trehalose and rhamnose were increased in *DUF581-9 OE* lines, while major carbohydrates like glucose or fructose were decreased. Overexpression of DUF581-9 seems to grossly affect amino acid levels, most of which show a decrease in the light as well as during the dark. Notably, *DUF581-9* overexpression plants show reduced arginine, glutamate, glutamic acid as well as GABA levels in the dark relative to control and *duf581-9* lines. In addition, intermediates of the tricarboxylic acid cycle such as citric acid, succinic acid, malate and fumaric acid were decreased in *DUF581-9 OE* lines particularly during the dark treatment.

These data thus suggest that *DUF581-9 OE* lines are affected in their ability to induce metabolic adjustments during dark induced starvation.

Given the previous observation that DUF581-9 interacts with KIN10/11 in yeast (Nietzsche *et al*., 2014), we assumed that the effect of DUF581-9 overexpression on SnRK1 signalling is directly mediated by this interaction. In order to test this hypothesis, the ectopically expressed DUF581-9 protein was pulled down from crude extracts by virtue of its C-terminal myc-tag. Subsequent immunoblotting of the eluate using an anti-KIN10 antibody yielded a strong signal in the pull-down from the transgenic lines while no signal was observed in the wild type control (Fig. S4a). This indicates that the ectopically expressed DUF581-9 protein interacts with KIN10 *in planta*. Furthermore, a mass spectrometric analysis of the eluate identified all SnRK1 subunits to be pulled down by DUF581-9 (Table 1). These data strongly suggest that DUF581-9 associates with the SnRK1 heterotrimeric complex *in planta*. No direct interaction of DUF581-9 with subunits of the SnRK1 complex other than KIN10 could be observed in yeast indicating that these are pulled down indirectly by virtue of their binding to the catalytic subunit (Fig. S4b).

**Table 1.** SnRK1 subunits identified in a DUF581-9-myc IP-MS.

| Gene name | Sum<br>PEP<br>Score | Coverage<br>[%] | # Peptides | # PSMs | Col-0 | DUF581-9 OE |
| --- | --- | --- | --- | --- | --- | --- |
| DUF581-9 (FLZ3) | 193,012 | 57 | 12 | 2733 | - | + |
| KIN10/KIN11 | 69,004 | 23 | 11 | 251 | - | + |
| KINβ1 | 12,888 | 21 | 3 | 54 | - | + |
| KINβ2 | 27,531 | 30 | 6 | 232 | - | + |
| KINβ $\gamma$ (SNF4) | 50,905 | 15 | 8 | 458 | - | + |

We next assessed the effect of DUF581-9 on SnRK1 activity in a protoplast transactivation assay based on the SnRK1 responsive promoter of the Arabidopsis *cyPPDK* gene fused to the GUS reporter gene (Henninger *et al*., 2022). Co-transfection of Arabidopsis protoplasts with the *pPPDK2::GUS* plasmid and a construct expressing KIN10 led to significant induction of the GUS reporter gene as compared reporter plasmid only control (Fig. 2d). When the same plasmid combination was supplemented by a DUF581-9 expressing construct, no significant induction of GUS activity could be observed.

Taken together, the data strongly suggest that DUF581-9 acts as a negative regulator of SnRK1 signalling in Arabidopsis through direct interaction with the SnRK1 holocomplex.

### DUF9 prevents activation of KIN10 by its upstream kinase GRIK

To further explore the inhibitory mechanism of DUF581-9 on KIN10, we employed an *in vitro* kinase assay based on *E. coli* produced recombinant proteins coupled to a sensitive and specific staining of phosphorylated proteins in SDS gels. The assay mixture contained GRIK2, which is required for T-loop phosphorylation (T175) of KIN10 and thus activation of the catalytic α subunit KIN10, and the SAMS peptide fused to MBP acting as a generic SnRK1 substrate. When recombinant GRIK2, KIN10 and SAMS were combined into the same assay mixture, phosphorylation signal for all three proteins could be detected (Fig. 3a). In contrast, the inclusion of recombinant DUF581-9 into the assay mixture strongly suppressed KIN10 phosphorylation, also preventing SAMS phosphorylation by KIN10 (Fig. 3a), indicating that DUF581-9 interferes with SnRK1 activity *in vitro*. Addition of the DUF581-9_C47S_ variant, previously demonstrated to be unable to bind KIN10 in yeast (Nietzsche *et al*., 2014), did barely reduce the KIN10 phosphorylation signal as well as SAMS phosphorylation *in vitro* (Fig. 3a). This indicates that the full inhibitory effect of DUF581-9 on KIN10 requires intactness of DUF581. In addition, DUF581-9 yields a phosphorylation signal in the presence of KIN10 and thus serves as a KIN10 phosphorylation substrate *in vitro* independent of the C47S substitution (Fig. 3a). When KIN10 was replaced by a catalytically inactive ATP-binding site mutant (KIN10_K48R_), no auto-phosphorylation of KIN10 as well as no DUF581-9 or SAMS signal was detected (Fig. 3a). To assess SAMS phosphorylation by KIN10 in a quantitative manner, we measured the incorporation of radiolabelled phosphate into the peptide using the same *in vitro* assay setup as above. The measurement revealed a significant reduction in the incorporation of label in the presence of DUF581-9 when compared to control conditions (Fig. 3b). Here, the presence of DUF581-9_C47S_ had no inhibitory effect on KIN10 activity but rather led to a stimulation of label incorporation likely through increased DUF581-9_C47S_ phosphorylation. Specificity of the assay conditions for KIN10 activity were verified by replacing the native enzyme with its inactive KIN10_K48R_ variant (Fig. 3b).

**Fig. 3.**
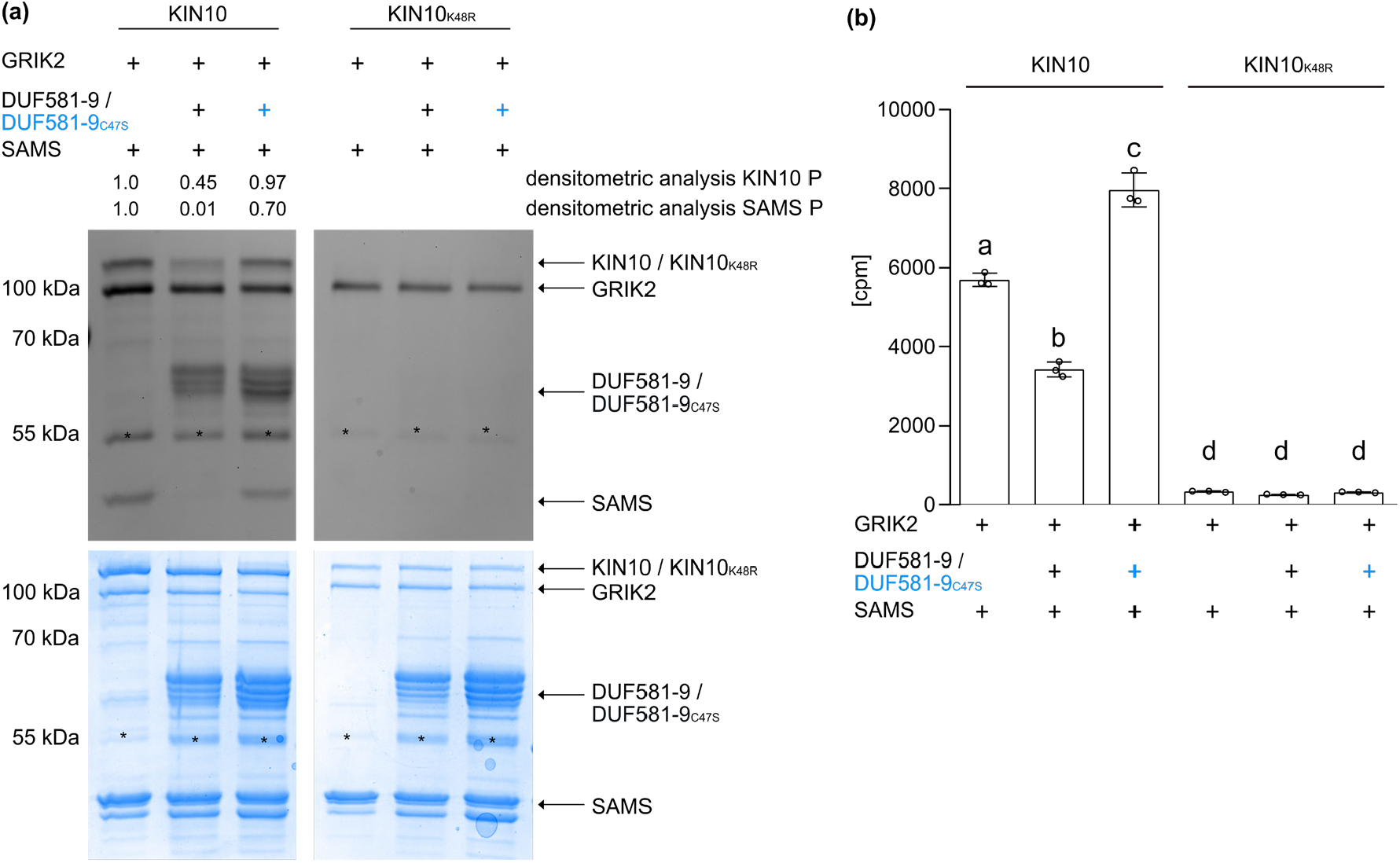
DUF581-9 inhibits KIN10 kinase activity *in vitro*. Analyses of KIN10 kinase activity *in vitro* using recombinant MBP-tagged full length GRIK2, KIN10, KIN10_K48R_, DUF581-9, DUF581-9_C47S_ and the generic KIN10 substrate SAMS. Protein phosphorylation was visualised by (a) ProQ Diamond Phosphostain^®^ including densitometric analysis of KIN10 and SAMS phosphorylation levels, as well as the respective Coomassie stained protein loading control. Asterisks indicate an unspecific protein band. Experiment was carried out at least three times with similar results. (b) *In vitro* kinase assay using ^32^P-γATP was performed with MBP-tagged recombinant proteins. Catalytically inactive MBP-KIN10_K48R_ served as a negative control. Error bars indicate SD. Asterisks mark statistical significant differences relative to the control without DUF581-9 or DUF581-9_C47S_ (ANOVA analysis followed by Tukey’s multiple comparison test). Three repetitions showed similar results.

An incremental reduction of DUF581-9 amount in the assay mixture led to a partial recovery of SAMS phosphorylation and thus KIN10 activity (Fig. 4a). A densitometrical analysis of SAMS phosphorylation revealed a strong negative correlation between the amount of DUF581-9 and SAMS staining intensity (Fig. 4b). However, no correlation was found between the DUF581-9 phosphorylation signal and the amount of SAMS present in mixture (Fig. S5). This indicates that the inhibitory effect of DUF581-9 on SAMS phosphorylation is not due to substrate competition.

**Fig. 4.**
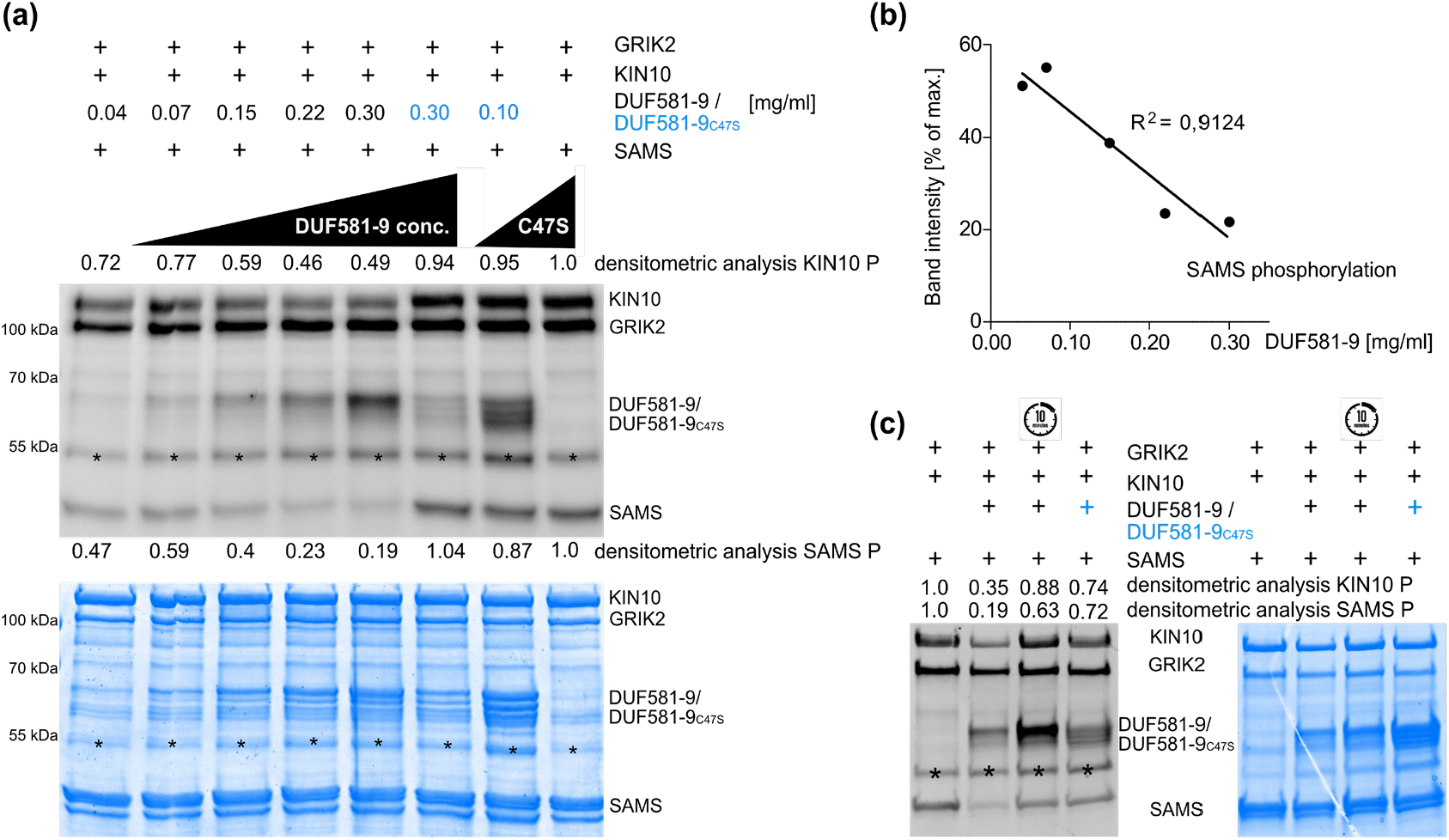
KIN10 inhibition by DUF581-9 is dose-dependent. Assay of *in vitro* KIN10 kinase (full length) activity depending on DUF581-9 amount. Protein phosphorylation levels were visualized by (a) ProQ Diamond Phosphostain® and quantified by densitometric analysis of phosphorylated KIN10, SAMS and DUF581-9/DUF581-9_C47S_ protein phosphorylation levels. One low and one high concentration was used for the DUF581-9_C47S_ negative control. As positive control served a kinase assay mixture containing GRIK2, KIN10 as well as its SAMS substrate (far right lane). (b) Negative correlation of SAMS phosphorylation levels and DUF581-9 protein amounts. (c) Kinase assay was performed as before, except that DUF581-9 was added to the mixture 10 min after all other components. Coomassie staining served as protein loading control. Asterisk indicates an unspecific protein band.

Since DUF581-9 has no obvious enzymatic function that could interfere with KIN10 phosphorylation, the dose-dependence of its inhibitory effect points towards a stoichiometric mechanism of KIN10 inhibition. Indeed, when GRIK2, KIN10 and SAMS were allowed to pre-incubate for 10 min before the addition of DUF581-9 the inhibitory effect on KIN10 and SAMS phosphorylation was abolished (Fig. 4c). This shows that DUF581-9 is not able to interfere with KIN10 activity once the kinase has been activated by GRIK2 or revert its activation, suggesting a steric mechanism of inhibition.

### DUF581-9 interferes with T-loop phosphorylation of KIN10

Phosphorylation of the T-loop within the catalytic domain of the SnRK1 α subunit is essential for its activation (Baena-González *et al*., 2007; Shen *et al*., 2009; Crozet *et al*., 2010; Rodrigues *et al*., 2013). To narrow down the effect of DUF581-9 on KIN10 phosphorylation, we assessed the T-loop phosphorylation *in vitro* more directly using an anti-phospho-α-AMPK (T172) antibody that specifically recognizes KIN10 phosphorylated at T175 (Sugden *et al*., 1999; Baena-González *et al*., 2007). T-loop phosphorylation of KIN10 by GRIK2 was readily detectable in the absence of DUF581-9 while no signal was observed upon inclusion of the DUF581 protein (Figure 5a). In turn, addition of the DUF581-9_C47S_ variant did not reduce the T-loop phosphorylation signal. In order to obtain a broader picture of the effect of DUF581-9 on KIN10 *in vitro* phosphorylation, specific sites were mapped using LC-MS/MS. When GRIK2, KIN10 and the DUF9_C47S_ variant were present in the assay mixture, four phosphorylation sites in addition to T175 could robustly be detected within the KIN10 polypeptide (S29; S338/339; S361) (Fig. 5b). As these sites remain un-phosphorylated in the KIN10_K48R_ variant, they likely result from auto-phosphorylation triggered by initial T-loop phosphorylation through GRIK2 (Fig. 5b). A quantitative analysis revealed that the presence of DUF581-9 significantly reduced protein phosphorylation at all sites up to approximately 70 % (Fig. 5b). Phosphorylation of GRIK2 activation loop (T153) was unaffected by DUF581-9, indicating that GRIK2 activity is not inhibited (Table S2).

**Fig. 5.**
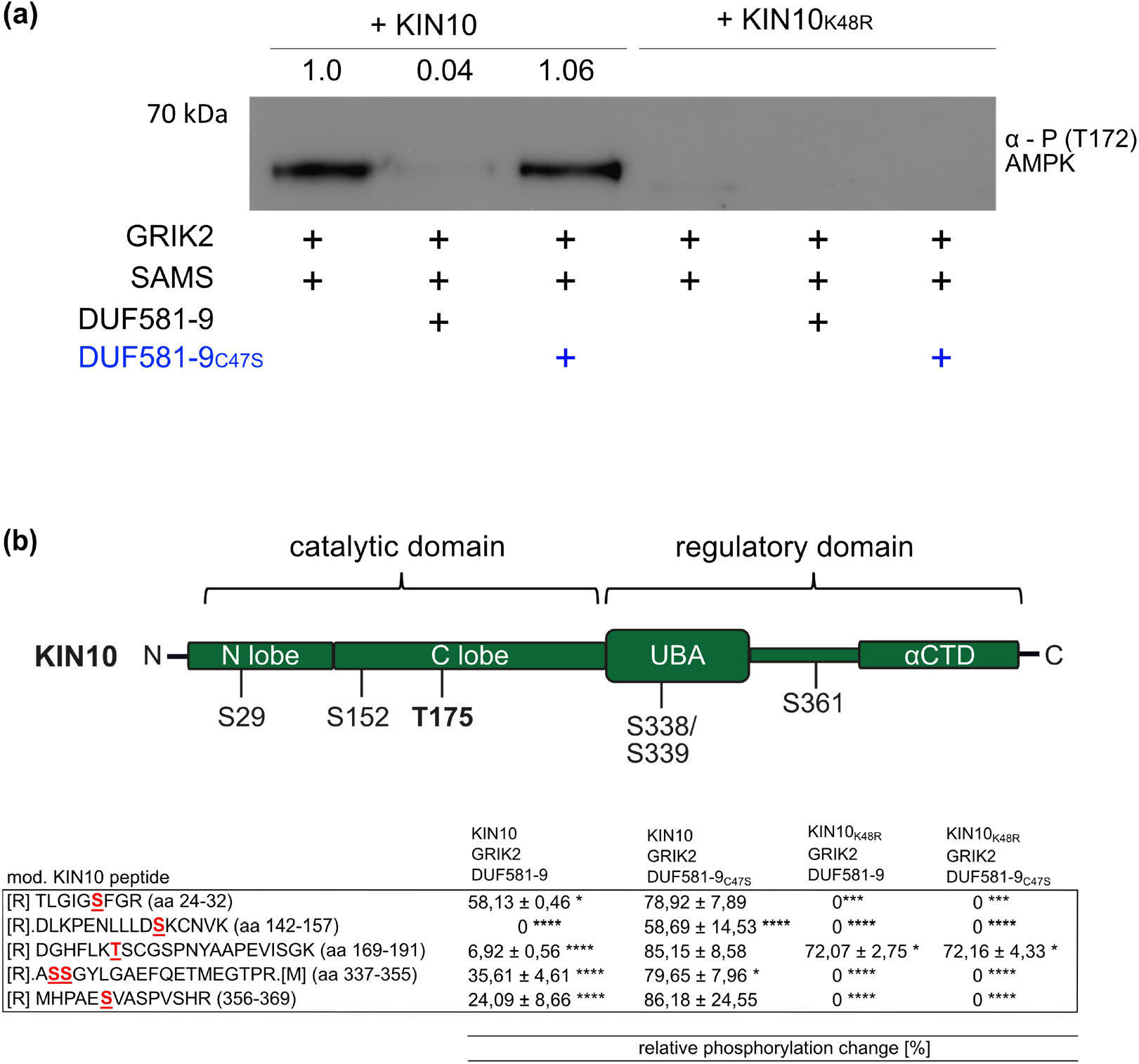
DUF581-9 interferes with KIN10 T175 T-loop phosphorylation. (a) Western blot analysis of recombinant KIN10 (full length) kinase assays using an anti-phospho-α-AMPK (T172) antibody specifically detecting phosphorylated KIN10 T175. (b) Mapping of KIN10 phosphorylation sites by LC-MS/MS. Schematic representation of robustly identified phosphorylation sites. Table indicates the relative changes in phosphorylation, calculated for different phosphorylation sites within the KIN10 polypeptide and compared to a positive control without DUF581-9 or DUF581-9_C47S_. Three technical replicates were measured and used for calculation. All experiments were carried out three times with similar results.

Thus, it is tempting to speculate that binding of DUF581-9 to KIN10 prevents the initial T-loop phosphorylation by GRIK2, which subsequently limits KIN10 activity and restricts auto-phosphorylation as well as downstream substrate phosphorylation.

### DUF581-9 binds to the KIN10 catalytic domain to weaken GRIK/KIN10 interaction

The above data suggest that DUF581-9 binds to KIN10 to somehow shield the T-loop from being phosphorylated by GRIK. This mechanism would require DUF581-9 to interact with KIN10 close to or at the region of either GRIK binding or phosphorylation, i.e. close to the T-loop. A direct Y2H assay demonstrates that the mere 290-amino acid N-terminal catalytic domain (CD) of KIN10 is necessary and sufficient for DUF581-9 binding while no interaction with the C-terminal regulatory domain could be detected (Fig. S6a). We also tested whether DUF581-9 could directly interact with GRIK2 in yeast; however, no binding between these two proteins could be observed (Fig. S6b). It has previously been shown that GRIK2 forms a stable complex with the CD of KIN10 *in vitro* (Shen *et al*., 2009). Because GRIK2 and DUF581-9 interact with the same region of KIN10, they may compete with each other for binding to KIN10. To test this hypothesis, we developed a BiFC-based assay to examine the KIN10 – GRIK2 dissociation by DUF581-9 *in vivo*. We co-infiltrated Agrobacterium strains carrying either a construct expressing KIN10-Venus^N173^ or GRIK2-Venus^C155^. Furthermore, a third construct expressing either DUF581-9-mCherry or a control mCherry fusion protein was co-infiltrated into leaves of *N. benthamiana*. Microscopic imaging and quantification of the fluorescence signal as a proxy for binding efficiency revealed that KIN10-Venus^N173^ and GRIK2-Venus^C155^ yielded readily detectable fluorescence (Figs. 6a and b), indicative for protein-protein interaction. The BiFC signal was significantly reduced in cells that additionally expressed DUF581-9-mcherry, suggesting a reduced interaction between KIN10 and GRIK2 (Figs. 6a and b). Co-expression of the fructose-1,6-bisphosphatase (FBPase)-mCherry control protein did not reduce BiFC fluorescence, although quantification of the mCherry fluorescence revealed comparable intensities between both combinations (Fig. 6b). In turn, DUF581-9-mcherry co-expression did not affect the BiFC of FBPase-Venus^N173^ or FBPase-Venus^C155^ homomerization although the mCherry fluorescence signal was in a similar range as before. Expression of all fusion proteins under investigation was verified by immunoblotting (Fig. S7). These results indicate that DUF581-9 specifically reduces the interaction between KIN10 and GRIK2 *in planta*.

**Fig. 6.**
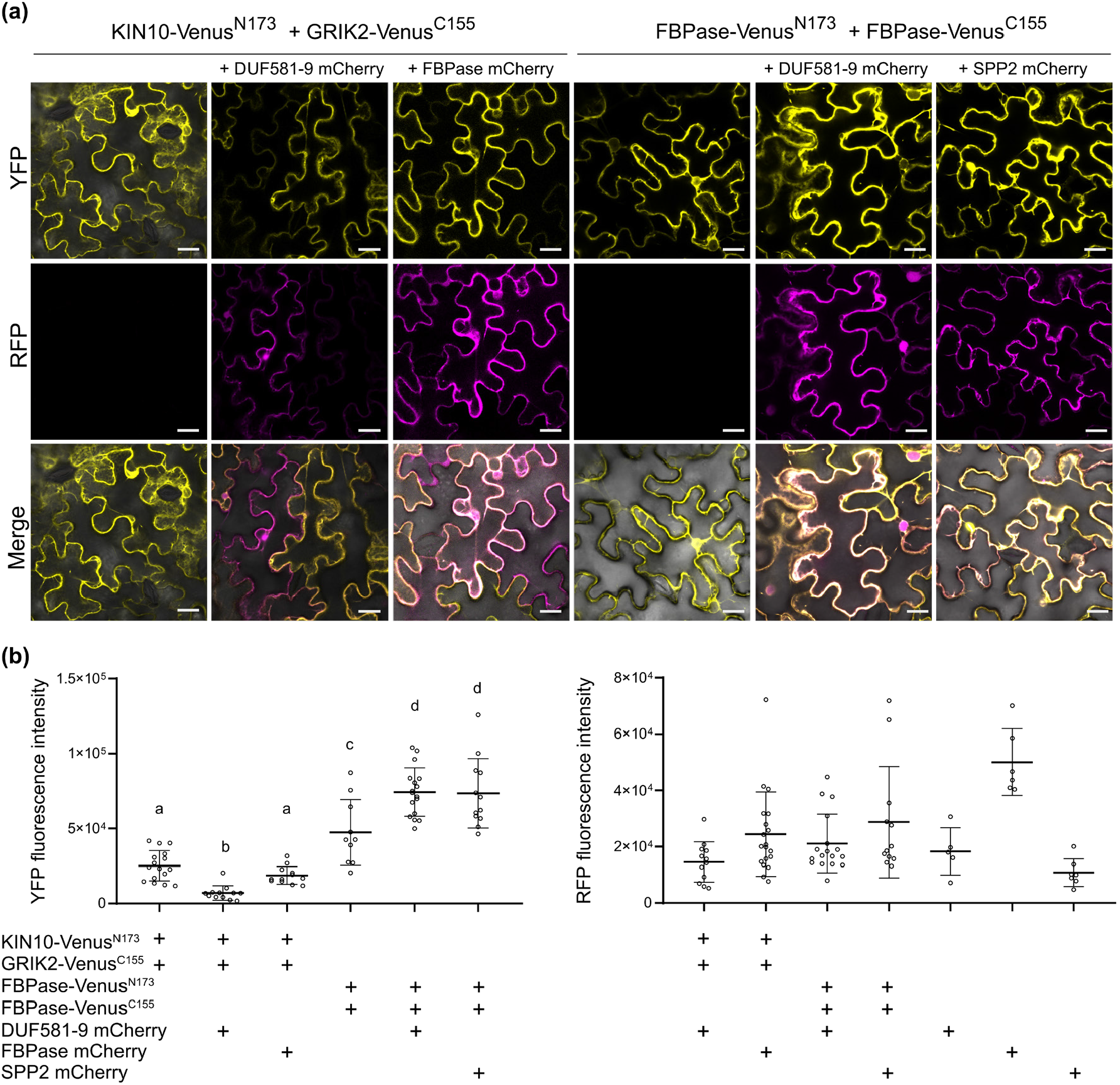
DUF581-9 weakens the interaction between KIN10 and GRIK2 *in vivo*. (a) DUF581-9 mCherry disrupts the BiFC signal generated from the association of KIN10-Venus^N173^ and GRIK2-Venus^C155^ *in planta*. Proteins were transiently expressed in leaves of *N. benthamiana* using *Agrobacterium*-infiltration and yellow fluorescent protein (YFP) and red fluorescence (mCherry) was monitored by confocal microscopy 48 hpi. Upper panel shows reconstituted YFP fluorescence in absence of a third partner. Middle panel shows mCherry fluorescence of either DUF581-9 mCherry or FBPase mCherry used as a control. Lower panel shows the merge of YFP and mCherry fluorescence. The homomerization of FBPase-Venus^N173^ and FBPase-Venus^C155^ was neither affected by DUF581-9 mCherry nor by SPP2 mCherry. The scale bar represents 20 µM. (b) The fluorescence signal intensity of YFP and mCherry (effector) were determined along a line drawn on the confocal images using ImageJ software. Letters above dots indicates statistically significant differences between the samples against KIN10-Venus^N173^ and GRIK2-Venus^C155^ control as determined by Oneway ANOVA followed by Tukey’s multiple comparisons test. The experiment was carried out twice with similar results.

An *in vitro* pull-down competition assay using recombinant proteins revealed that GST-KIN10 was able to pull-down either MBP-GRIK2 or MBP-DUF581-9 when each protein was added alone (Fig. 7). The addition of free MBP as a third protein did not affect the ability of GST-KIN10 to pull-down MBP-GRIK2. However, the interaction was substantially weakened when MBP-DUF581-9 was present as a third protein (Fig. 7). Although the MPB-DUF581-9_C47S_ variant was still able to bind KIN10 *in vitro* it had no obvious effect on the ability of GST-KIN10 to pull-down MBP-GRIK2 (Fig. 7).

**Fig. 7.**
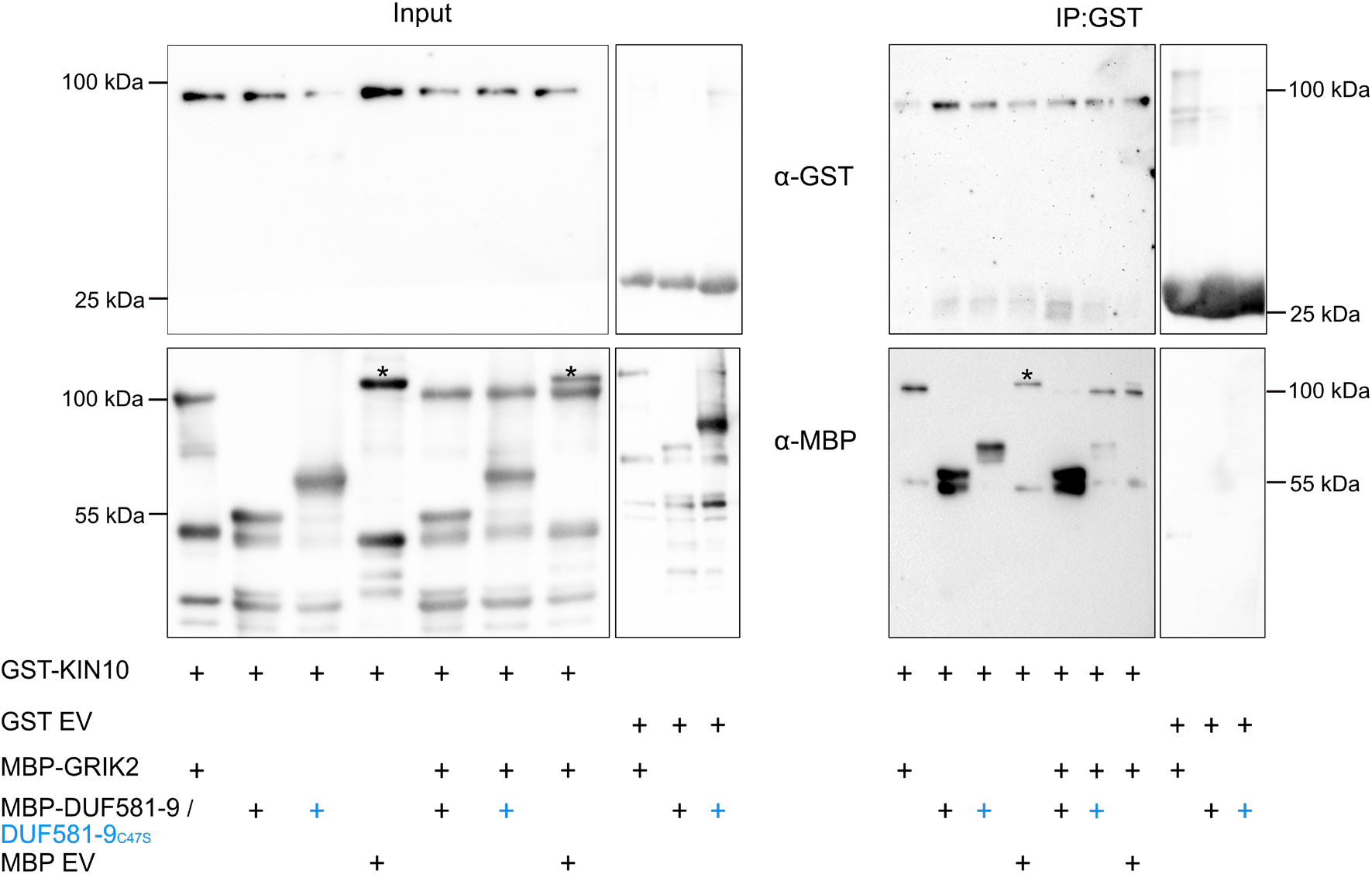
DUF581-9 interferes with KIN10/GRIK2 interaction *in vitro*. Proteins were mixed as indicated and KIN10-GST (full length) was pulled down from the mixture. Recombinant proteins were detected before (Input) and after (IP:GST) by immunoblotting using anti-MBP or anti-GST antibodies. The experiment was carried out at least twice with similar results. Asterisk indicates an unspecific protein band.

Taken together, the data suggest that DUF581-9 prevents KIN10 activation by interfering with GRIK binding and eventually blocks phosphorylation of the critical T-loop.

### DUF581-9 undergoes proteasomal degradation under SnRK1 activating conditions

The question arises how KIN10 complexed with DUF581-9 can be activated when SnRK1 signalling is required. One possible mechanism could include removal of KIN10 bound DUF581-9 by proteolysis. Therefore, we monitored DUF581-9 protein stability in CaMV35S-driven overexpression lines under energy limiting conditions. A DUF581-9-myc protein signal is readily detectable in samples taken from illuminated plants while the same signal vanished when the plants were subjected to dark-induced starvation for 6 h (Fig. 8). However, when the proteasome inhibitor bortezomib was infiltrated before initiation of starvation, DUF581-9 protein degradation was greatly diminished. Analysis by qRT-PCR of *DUF581-9* expression excluded an effect of the treatment on mRNA levels (Fig. S8), indicating destabilisation and degradation of the protein under energy limiting conditions by the proteasome.

**Fig. 8.**
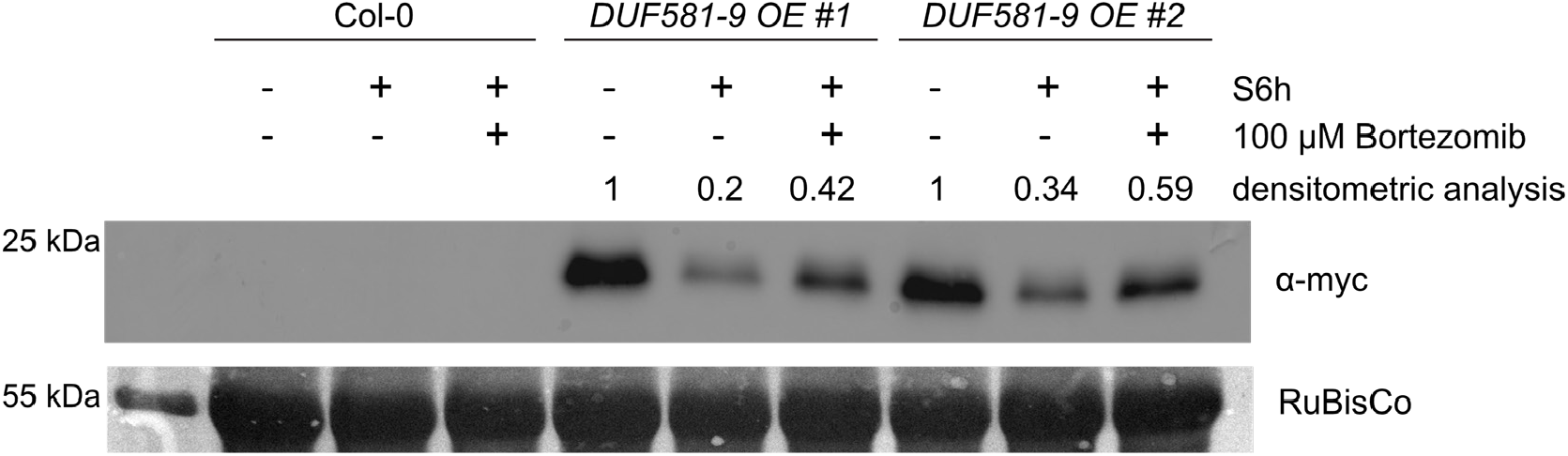
Ectopically expressed DUF581-9 is degraded under dark-induced starvation. 4 week old Col-0 and *DUF581-9 OE* plants were subjected to 6 h darkness (S6h) whereas control plants were sampled during the light phase. Myc affinity trap was added to plant raw extract to enrich myc-tagged DUF581-9 protein followed by immunoblot analysis using an α-myc antibody. Densitometric analysis reflects changes in protein levels between light and dark treated plants. The experiment was repeated three times with similar results.

## Discussion

SnRK1 protein kinases are central regulators of energy and stress signalling in plants integrating a multitude of environmental stimuli into a range of metabolic and developmental responses (Crepin &Rolland, 2019). SnRK1 activity is under complex regulation involving post-translational modifications, metabolites, protein-protein interactions and complex localization (Crozet *et al*., 2014; Broeckx *et al*., 2016). However, how SnRK1 controls and coordinates this diversity of processes in time and space to ensure a proper signalling response in a context specific manner, is largely unknown.

In this study, we provide evidence that the DUF581-9 protein acts as a negative regulator of SnRK1 activity, which is in line with a recent hypothesis that SnRK1 is activated by default but its activity is repressed under conditions when signalling is not required (Crepin &Rolland, 2019; Ramon *et al*., 2019). Consistent with a role as negative regulator of SnRK1 activity, DUF581-9 expression is reduced during dark-induced starvation but induced during the recovery phase, likely to rapidly shut down starvation signalling when conditions ameliorate. DUF581-9 expression does not respond to hypoxia, although this also leads to cellular energy depletion and concomitant activation of SnRK1 signalling (Cho *et al*., 2012). This opens the possibility that DUF581-9 is only involved in SnRK1 regulation under certain types of stresses.

Evidence for a role of DUF581-9 as negative regulator of SnRK1 signalling *in vivo* was obtained through the analysis of transgenic Arabidopsis lines with altered *DUF581-9* expression. Constitutive overexpression of DUF581-9 decreased the expression of SnRK1 marker genes during short-term dark induced starvation and also reduced plant survival after long-term starvation treatment. Dark treatment leads to a rapid decline in cellular energy levels resulting in the induction of alternative pathways to generate ATP from non-carbohydrate resources such as proteins, fatty acids or chlorophyll catabolism (Usadel *et al*., 2008; Araujo *et al*., 2011; Zhu *et al*., 2022). This response is at least partially dependent on SnRK1 mediated transcriptional regulation (Baena-González *et al*., 2007; Pedrotti *et al*., 2018). Arabidopsis plants overexpressing *DUF581-9* show distinct metabolic changes relative to wild type that become even more pronounced during dark-induced starvation and are diagnostic for impaired SnRK1 signalling. An early response to dark-induced starvation is the accumulation of amino acids deriving from protein degradation (Zhu *et al*., 2022), which is generally reduced in *DUF581-9* overexpression lines. The contents of metabolites related to N-recycling as a result from protein degradation (Kennedy *et al*., 2019) such as glutamate, glutamine, arginine and urea is reduced in dark-treated *DUF581-9* overexpression lines as compared to the control, which is in line with a reduced amino acid accumulation. Energy production through the tricarboxylic acid (TCA) cycle fuelled by the breakdown of protein, lipids and chlorophyll is of critical importance to maintain high respiratory rates during starvation (Araujo *et al*., 2011). Several TCA intermediates such as succinate, citric acid, fumaric acid and malic acid are reduced in overexpression lines as compared to the wild type or *duf581-9* lines.

Taken together, the metabolite profiles of *DUF581-9* OE lines during early dark induced starvation strongly argue for an impairment of metabolic adaptation upon sudden energy limitation in these plants that can well be explained by a compromised SnRK1 signalling.

DUF581-9 was able to pull-down the entire SnRK1 complex from plant extracts, including the catalytic α subunit, the β subunits as well as the β/γ subunit. Other DUF581 isoforms have been shown to also interact with regulatory SnRK1 subunits in yeast (Carianopol *et al*., 2020); however, for DUF581-9 a direct interaction could only be observed with the catalytic subunit. This suggests, that DUF581-9 functions through direct binding to the readily assembled SnRK1 complex by a direct interaction with the catalytic α subunit. In line with a compromised SnRK1 response in *DUF581-9* overexpression lines, co-expression of KIN10 with DUF581-9 in protoplasts reduces the activation of a SnRK1-responsive reporter gene.

Thus, the data obtained so far suggest that DUF581-9 negatively regulates SnRK1 signalling through a direct interaction with the catalytic subunit of the kinase holocomplex.

The addition of DUF581-9 also inhibited KIN10 kinase activity in an *in vitro* reconstituted GRIK2/KIN10/SAMS cascade utilising *E. coli* produced recombinant proteins. This effect was dependent on an intact DUF581 domain, as the DUF581-9_C47S_ variant did not inhibit phosphorylation activity *in vitro*. Although a strong negative correlation between the concentration of DUF581-9 in the assay mix and the SAMS phosphorylation signal was observed, increasing concentrations of SAMS had no effect on the concentration of DUF581-9 phosphorylation. This suggests that KIN10 inhibition by DUF581-9 is not a result of both proteins competing for binding at the same site. Furthermore, DUF581-9 was not able to inhibit the activity of pre-activated KIN10, indicating it is not able to revert activation once it has occurred. These observations can be explained by a scenario in which DUF581-9 stoichiometrically binds to KIN10 to prevent its activation by steric or structural hindrance rather than through an inherent enzymatic activity.

The proposed sequence of events leading to KIN10 activation in the *in vitro* assay is as follows: (i) T153 auto-phosphorylation/activation of GRIK2; (ii) T175 KIN10 phosphorylation/activation by auto-activated GRIK2; (iii) reciprocal S260 GRIK2 phosphorylation/inhibition by activated KIN10 (Crozet *et al*., 2010). An effect of DUF581-9 on the activity of the up-stream activating kinase GRIK2 can largely be excluded since the GRIK2 auto-phosphorylation at T153, as a proxy for its activity, was not affected in the presence of DUF581-9. However, addition of DUF581-9 substantially reduced the KIN10 phosphorylation signal in phospho-stainings, correlating with a reduction in T-loop phosphorylation as revealed by immunoblotting using phosphorylation-site specific antibodies as well as by a quantitative assessment of KIN10 T-loop phosphorylation using LC-MS/MS. The presence of DUF581-9 also affected the phosphorylation level of additional sites within the KIN10 polypeptide. Phosphorylation of residues other than T175 is likely the result of KIN10 auto-phosphorylation as the catalytically inactive KIN10_K48R_ variant still shows detectable T175 phosphorylation, while additional sites remain un-phosphorylated. The presence of DUF581-9 also reduces KIN10 mediated GRIK2 S260 transphosphorylation (Crozet *et al*., 2010), further supporting an inhibitory effect of the protein on kinase activity. Whether the auto-phosphorylation sites detected *in vitro* correspond to the KIN10 auto-phosphorylation pattern *in planta* (Baena-González *et al*., 2007) and whether this is required for full kinase activity is currently unknown. Taken together, the data strongly suggest that DUF581-9 inhibits KIN10 activity by interference with GRIK2 mediated phosphorylation/activation, independent of affecting GRIK2 activity in general.

As prerequisite for T-loop phosphorylation GRIKs tightly interact with the SnRK1 catalytic α subunits (Shen *et al*., 2009). We show here that DUF581-9 weakens the interaction between GRIK2 and KIN10 *in vivo* and *in vitro*, likely with repercussions on T-loop phosphorylation efficiency. The sugar-signalling molecule T6P has been reported to inhibit SnRK1 activity in different plants (Zhang *et al*., 2009; Debast *et al*., 2011) and recent evidence suggests that similar to DUF581-9, T6P directly binds to the catalytic domain of KIN10 to weaken the interaction with upstream GRIK and thereby reducing KIN10 T-loop phosphorylation (Zhai *et al*., 2018). The mechanism through which T6P weakens the interaction between KIN10 and GRIK is currently unclear. For DUF581-9 it is possible that it competes with GRIK for the same binding site on the KIN10 surface and thereby reducing the accessibility of the T-loop for GRIK phosphorylation. Alternatively, DUF581-9 binding could induce structural changes in KIN10 that weaken GRIK binding.

Given the multitude of SnRK1 regulatory mechanisms that have already been described (Broeckx *et al*., 2016; Crepin &Rolland, 2019), the question remains what the biological significance of the regulation of SnRK1 signalling by DUF581-9 could be? Although *DUF581-9* expression was readily detectable in mature leaves by qRT-PCR, the DUF581-9 promoter GUS analyses suggest that expression mainly overlaps with the shoot apical meristem and with mitotic regions of developing leaves (Kazama *et al*., 2010), as well as in certain parts of the flower. This might also explain why *duf581-9* mutants display indeed enhanced expression of the SnRK1 marker gene *DIN6* during early starvation, but behave like wild type control plants with respect to metabolite changes and survival of long term starvation. The expression pattern of *DUF581-9* largely resembles that of GRIK1 and GRIK2 for which RNA was detectable at comparable levels in all tissues investigated but the protein was only found in the shoot apical meristem and very young leaves (Shen &Hanley-Bowdoin, 2006). It has been suggested that at least in proliferating young tissues, the upstream kinases are required for initial phosphorylation and activation of newly synthesized SnRK1 (Shen *et al*., 2009). A knock-out of both GRIK isoforms in Arabidopsis severely affected plant growth and development and double mutant plants did not grow without sugar supplementation (Glab *et al*., 2017). The double mutant contained wild-type levels of KIN10, but the protein exhibited no or highly reduced T-loop T175 phosphorylation with a concomitant reduction in catalytic activity (Glab *et al*., 2017). This suggests that SnRK1 activation through GRIK phosphorylation is essential for SnRK1 function and also for overall plant growth and development. Young proliferating tissues are likely to have a high energy demand and carbohydrate supply in form of sucrose and its metabolites have been shown to be a major regulator of cell cycle progression (Riou-Khamlichi *et al*., 2000). In addition to SnRK1 regulation through mechanisms such as regulatory β subunit myristoylation and nuclear exclusion of the pre-activated complex (Ramon *et al*., 2019), these cells might require an additional layer of SnRK1 regulation that safe-guards its unintentional activation under conditions prioritizing cell division and growth. Binding of DUF581-9 to the SnRK1α subunit would prevent the phosphorylation and thus pre-activation of newly synthesized KIN protein. Upon starvation conditions, DUF581-9 is rapidly turned over by the proteasome exposing the T-loop for phosphorylation by the constitutively active GRIK up-stream kinases and thus allowing for rapid SnRK1 activation without the need of de-novo KIN protein synthesis or complex assembly.

Future studies will have to clarify whether other members of the DUF581/FLZ protein family have similar functions in the regulation of SnRK1 activity and what role the additional interaction partners of these regulatory proteins play within the SnRK1 signalling network (Nietzsche *et al*., 2016).

## Acknowledgements

We thank Kerstin Bieler and Mandy Heinze for their skillful technical help. S.A. and A.R.F. acknowledge funding of the PlantaSYST project by the European Union’s Horizon 2020 Research and Innovation Programme (SGA-CSA no. 664621 and no. 739582 under FPA no. 664620).

## Author contributions

J.B. and F.B. conceived the study. J.B., J.L., S.A. and C.W. performed the experiments. J.B., J.L., S.A., C.W., A.R.F., W. D.-L. and F.B. analysed the data. J.B. generated the figures. J.B. and F.B. wrote the paper with input from all authors.

## Data availability

The data that support the findings of this study are available from the corresponding author upon request.

## Supporting Information

**Fig. S1.**
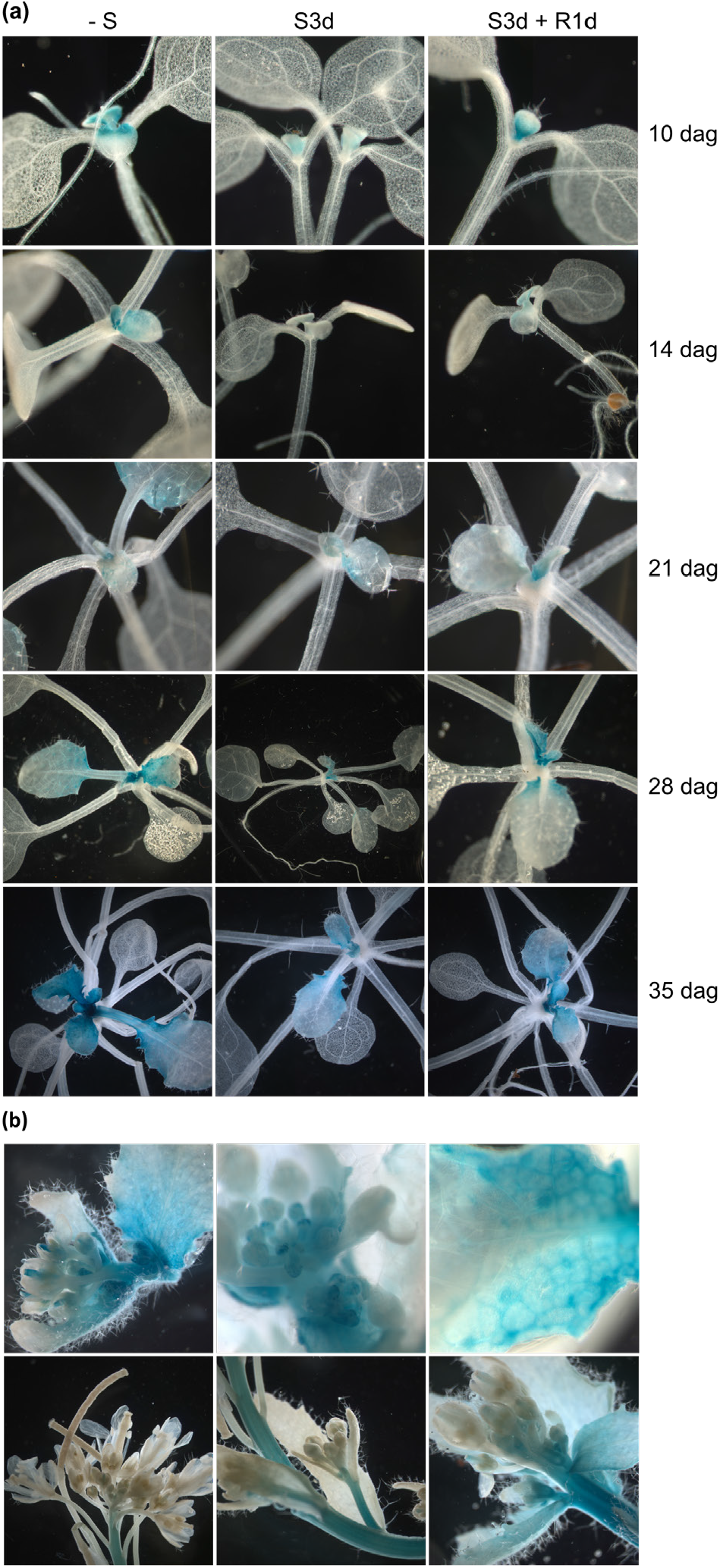
Tissue- and developmental stage-specific expression pattern of *DUF581-9* as revealed by the analysis of *Pro*^*DUF581-9*^*::GUS* Arabidopsis plants. (a) Shoots from plants of indicated ages (dag = day after germination) grown under short-day conditions (8h light/16h darkness) were stained for GUS activity. –S = plants grown in standard light / dark conditions; S3d = plants subjected to 3 days of dark treatment; S3d+R1d = plants analysed after one day of dark-treatment recovery in short day conditions. (b) GUS-histochemical staining of inflorescence tissue from plants grown in LD.

**Fig. S2.**
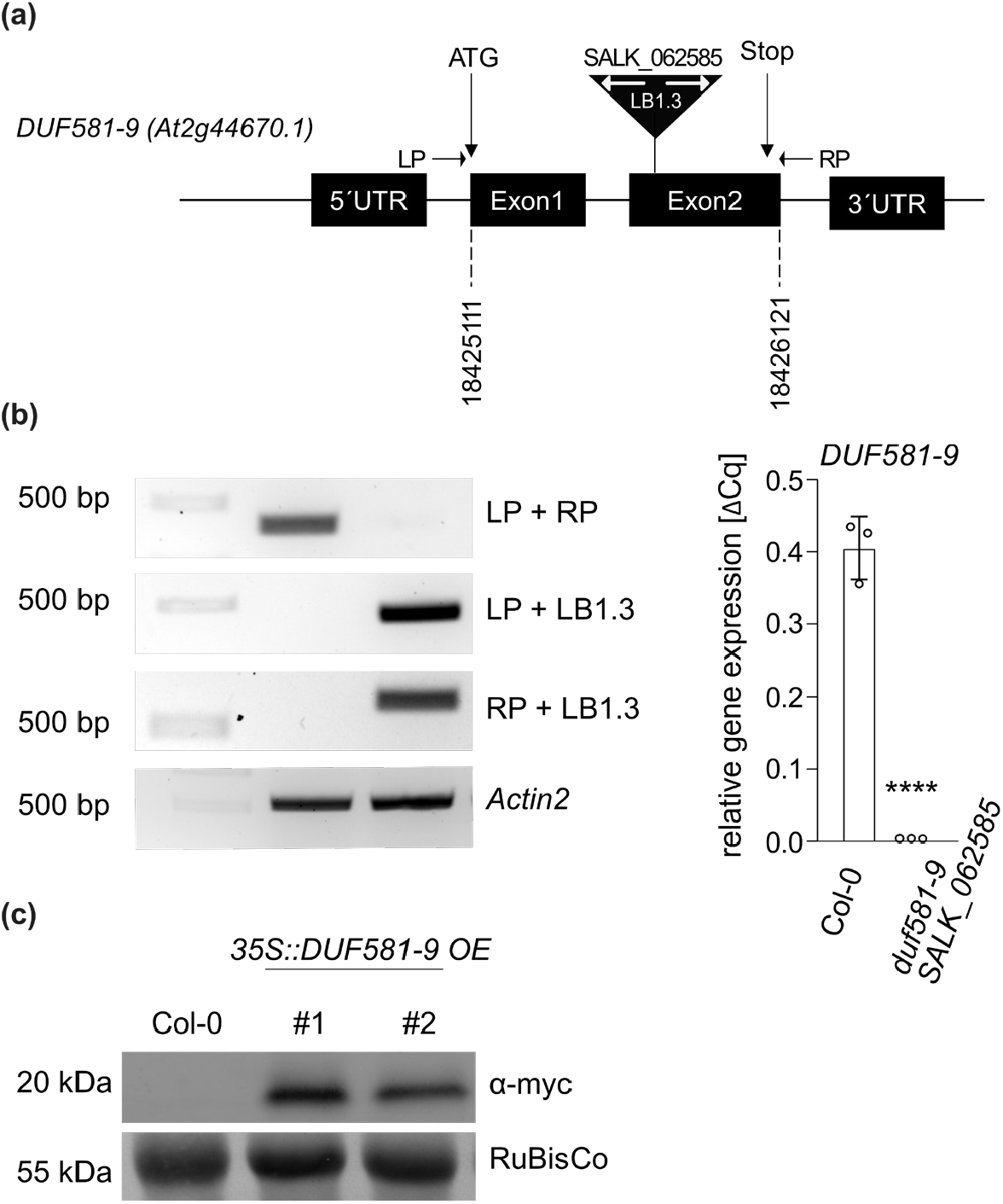
Characterization of Arabidopsis plants with modulated *DUF581-9* expression. (a) Schematic representation of the DUF581-9 (At2g44670) gene locus in *duf581-9* knock-out plants (SALK_062585). The T-DNA insertion sites and primer (LP, RP, and LB1.3) locations are indicated. (b) Genotyping of *duf581-9* plants by PCR on genomic DNA (left). The wild type *DUF581-9* allele is amplified with primers LP and RP while primer combinations LP/LB1.3 and RP/LB1.3 amplify a fragment only in the T-DNA insertion line. qRT-PCR analysis of *DUF581-9* expression in Col-0 and *duf581-9* knock-out plants (right). *UBC9* was used as reference gene. Error bars indicate SD. Asterisks indicated statistical significant differences relative to the Col-0 control (ANOVA analysis followed by Dunnett’s multiple comparison test) \*\*\*\**P* < 0.0001. (c) Immunoblot analysis of *35S::DUF581-9 OE* lines using an anti myc-antibody. Leaf material from four week old plants of the T3 generation was used for the analysis. Amido black stain of RuBisCo on the membrane served as loading control.

**Fig. S3.**
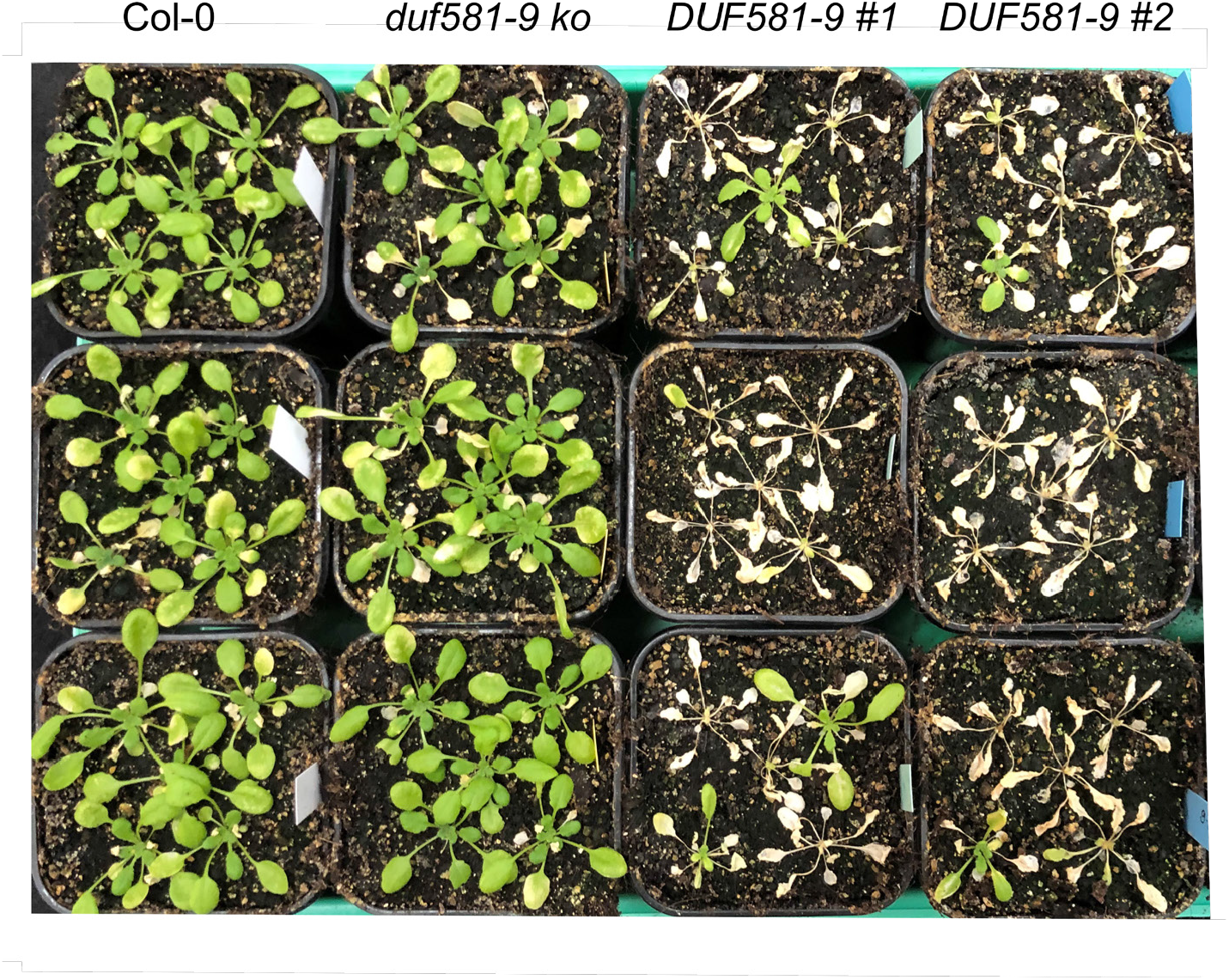
Overexpression of DUF581-9 reduces plant survival after long-term dark treatment. Long term starvation was conducted with 3 week old Arabidopsis plants grown under SD conditions. Plants were subjected to a dark-treatment for 10 days followed by a recovery phase of 7 days under normal light and growth conditions. Plants with a clearly visible intact green meristem were considered survivors.

**Fig. S4.**
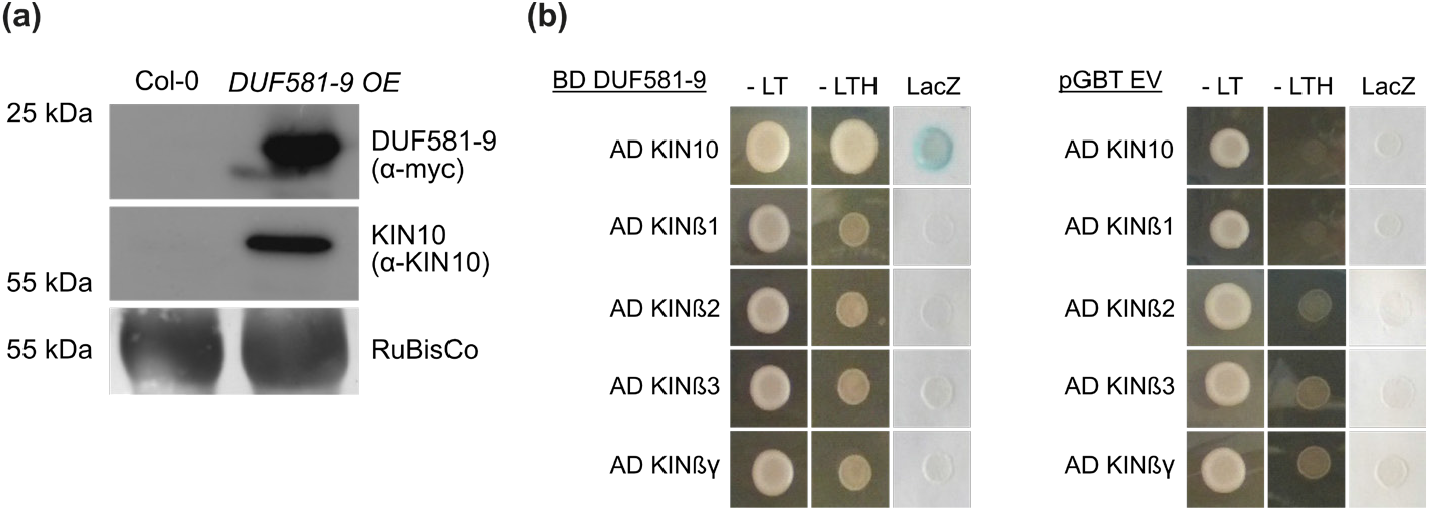
(a) Pull-down of DUF581-9 myc from transgenic Arabidopsis lines. DUF581-9 myc was pulled down from raw extracts prepared from *35S::DUF581-9 OE* plants using myc-Trap magnetic beads. An aliquot of the precipitate was used for immunoblotting with an α-myc antibody and α-KIN10 antibody. The experiment was carried out at least three times. (b) Yeast two-hybrid assay of DUF581-9 interaction with subunits of the Arabidopsis SnRK1 complex. Cells were grown on selective medium before a *lacZ* filter assay was performed.–LT, Yeast growth on medium without Leu and Trp; –HTL, yeast growth on medium lacking His, Leu, and Trp, indicating expression of the *HIS3* reporter gene; LacZ, activity of the *lacZ* reporter gene. Left, interaction of DUF581-9 fused to the GAL4 DNA binding domain (BD) with the different SnRK1 subunits fused to the GAL4 activation domain (AD). Right, test for interaction of AD constructs against the empty BD vector control. The experiment was accomplished twice with similar results.

**Fig. S5.**
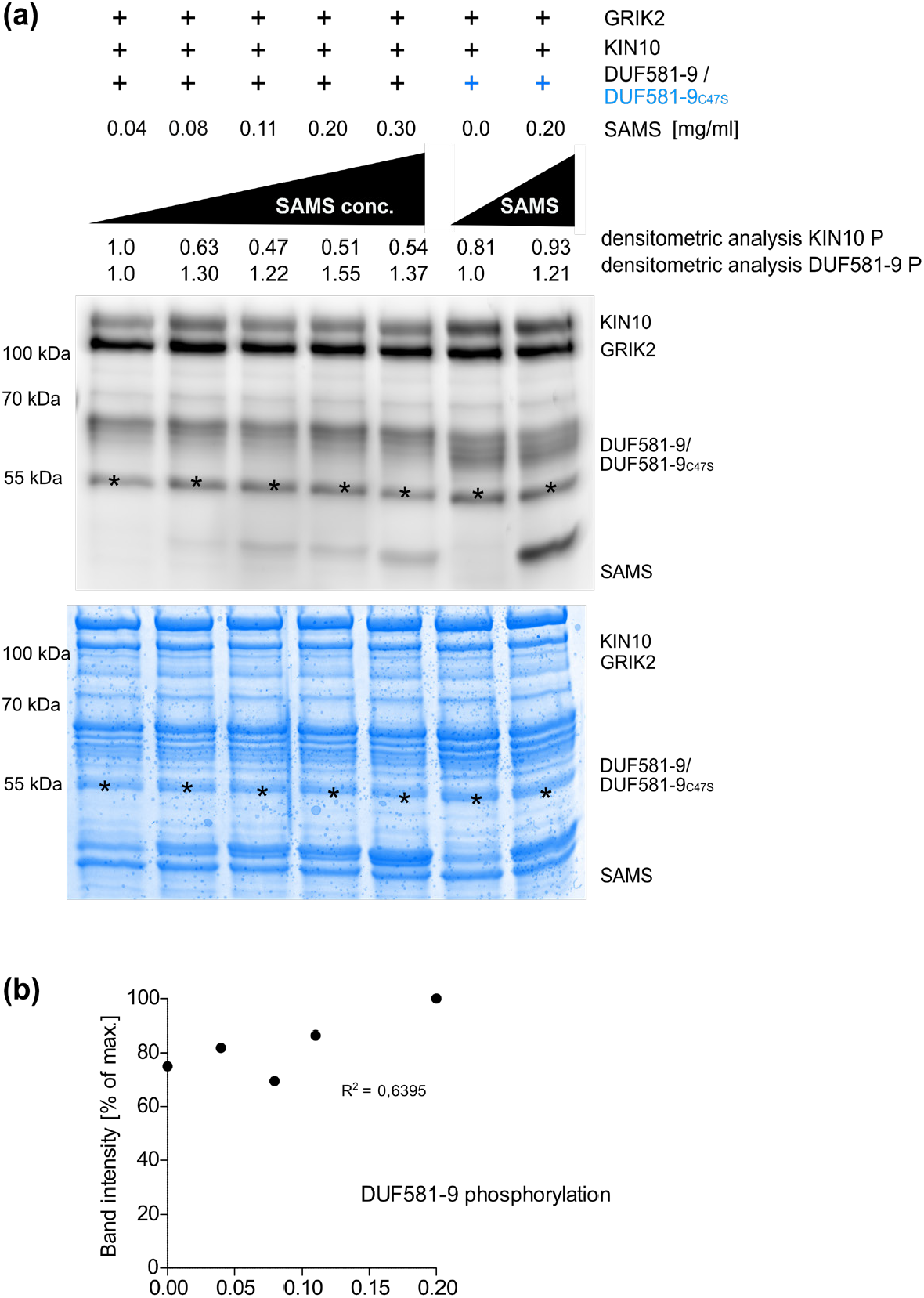
Increasing amounts of SAMS do not affect DUF581-9 phosphorylation. (a) *In vitro* SnRK1 kinase assay and ProQ Diamond Phosphostain® with densitometric analysis of phosphorylated DUF581-9 in the presence of increasing amounts of SAMS. Coomassie staining served as protein loading control. Asterisk indicates an unspecific protein band. (b) Correlation analysis of the DUF581-9 phosphorylation signal relative to the SAMS amount.

**Fig. S6.**
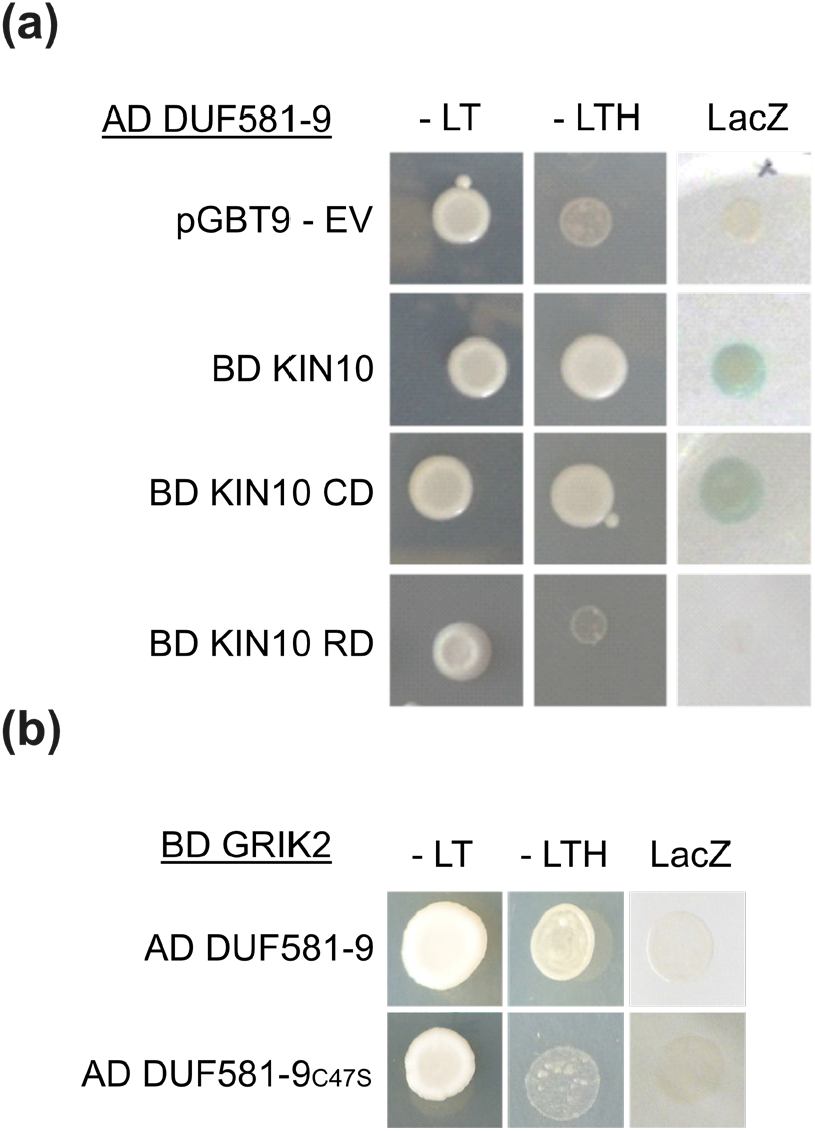
DUF581-9 interacts with the KIN10 catalytic domain but not with GRIK2. (a) Yeast two-hybrid assay of DUF581-9 interaction with either the KIN10 full-length protein (FL), the KIN10 catalytic domain (CD, aa 1 – 272), or the KIN10 regulatory domain (RD, aa 294 - 512). Cells were grown on selective medium before a *lacZ* filter assay was performed.–LT, Yeast growth on medium without Leu and Trp; –HTL, yeast growth on medium lacking His, Leu, and Trp, indicating expression of the *HIS3* reporter gene; LacZ, activity of the *lacZ* reporter gene. DUF581-9 fused to the GAL4 activation domain (AD), while KIN10 variants were fused to the GAL4 DNA binding domain (BD). pBD-EV = empty vector control. (b) DUF581-9 does not interact with GRIK2 in yeast.

**Fig. S7.**
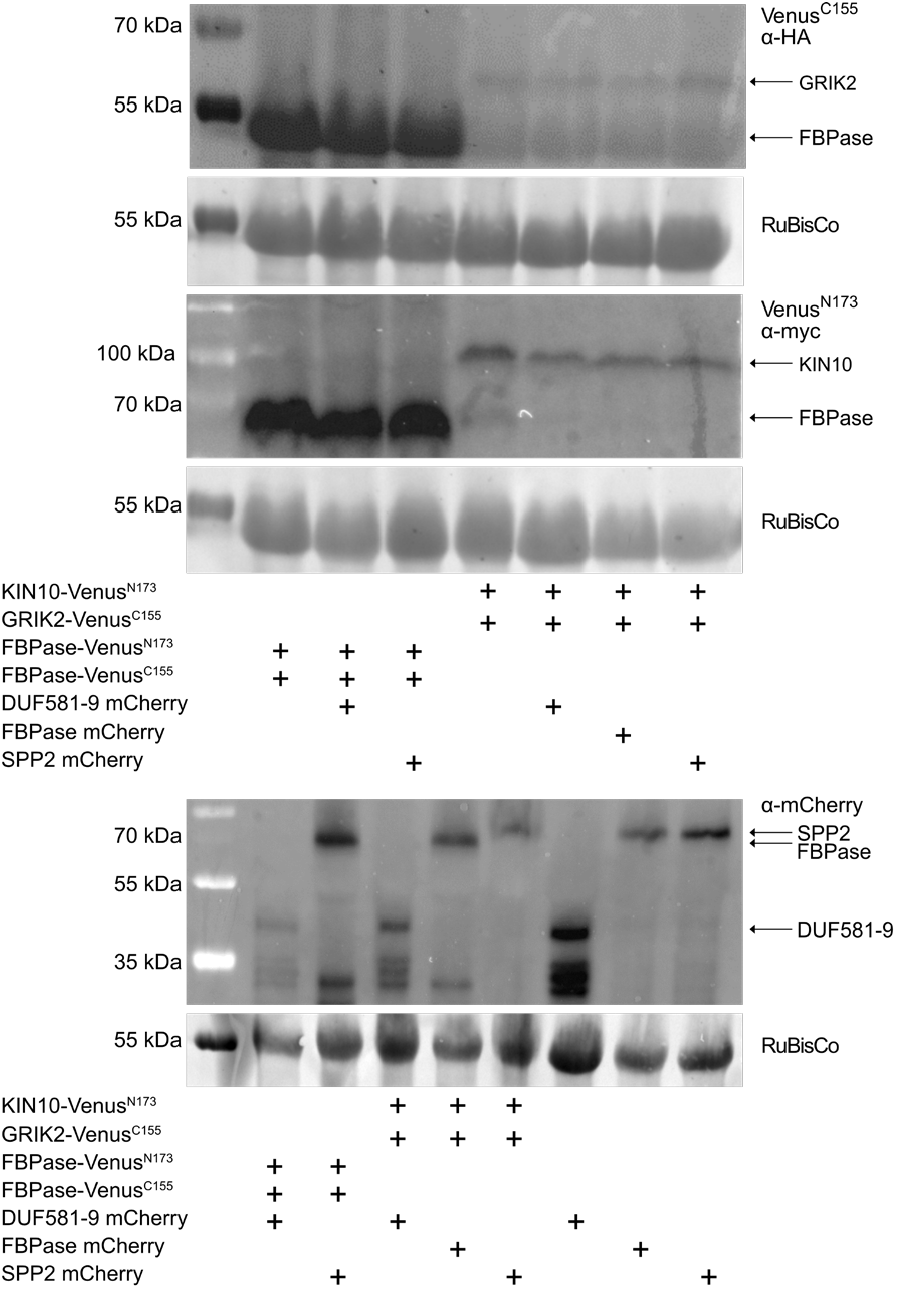
Immunoblot verification of protein expression in BiFC experiments. Immunoblot analysis of transiently expressed mCherry proteins as well as BiFC Venus^N173^ (α-myc) and BiFC Venus^C155^ (α-HA) constructs. Molecular weight of DUF581-9 (10,5 kDa), KIN10 (58,4 kDa), GRIK2 (45,7 kDa), FBPase (45,2 kDa), SPP2 (47,9 kDa) is gained by fusion to mCherry, Venus^N173^ or Venus^C155^ tag (27 kDa, 19,55 kDa and 7,3 kDa, respectively). Upper panel shows α-myc immunoblot, middle panel shows α-Ha and lower panel shows α-mCherry immunoblot. Amido black stain of RuBisCo on the membrane served as loading control.

**Fig. S8.**
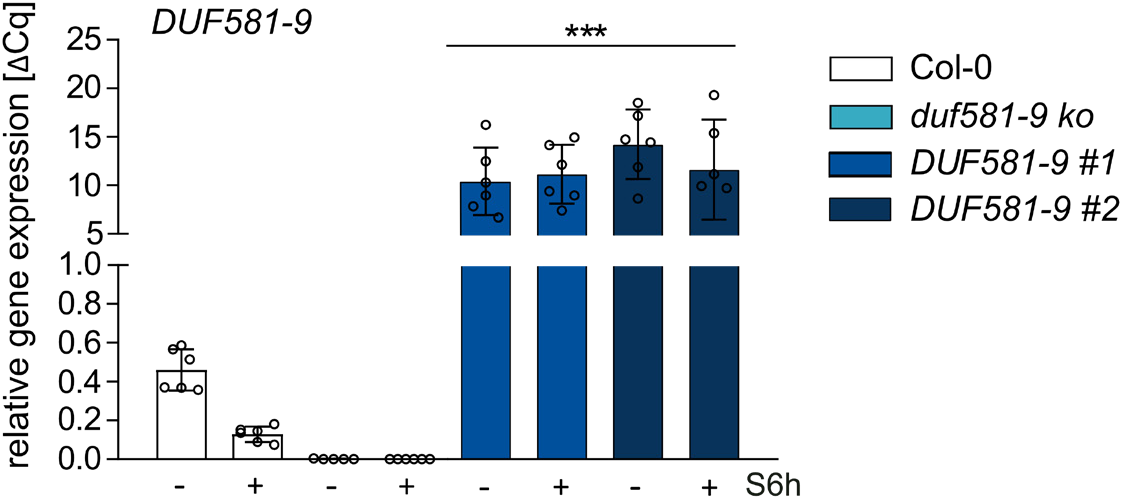
Starvation treatment does not affect DUF581-9 transcript levels in *35S::DUF581-9 OE* lines. Plants of *35S::DUF581-9 OE* line alongside with Col-0 wild type plants were subjected to 6 h of dark-induced starvation (S6h). Subsequently *DUF581-9* transcript levels were monitored using qRT-PCR. *UBC9* served as a control. Error bars indicate SD. Asterisks indicated statistical significant differences relative to the Col-0 control (ANOVA analysis followed by Sidak’s multiple comparison test) \*\*\**P* < 0.001. Experiment was repeated at least three times.

## Supplementary Material

Table S1. Metabolite levels

**Table S2.**
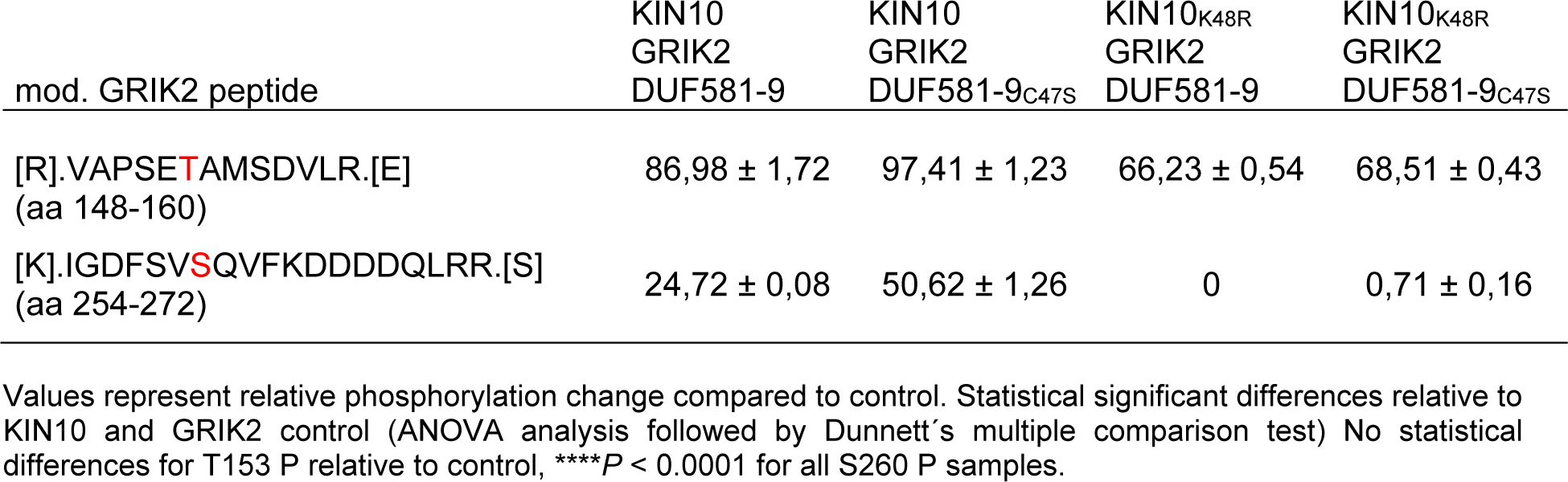
DUF581-9 does not interfere with phosphorylation of the GRIK2 activation loop.

**Table S3.**
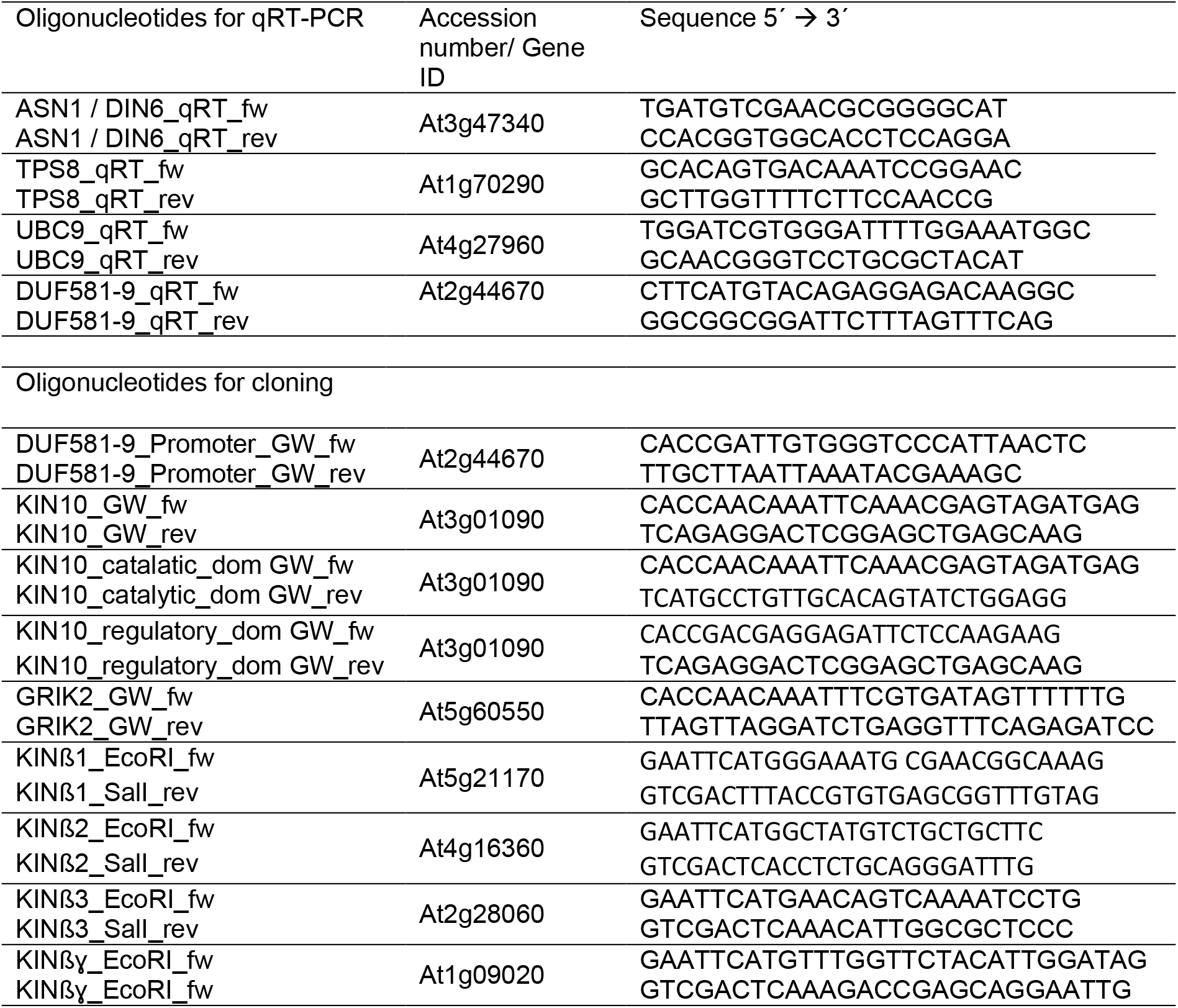
Oligonucleotides

